# TMEM65-dependent Ca^2+^ extrusion safeguards mitochondrial homeostasis

**DOI:** 10.1101/2023.10.10.561661

**Authors:** Massimo Vetralla, Lena Wischhof, Vanessa Cadenelli, Enzo Scifo, Dan Ehninger, Rosario Rizzuto, Daniele Bano, Diego De Stefani

## Abstract

The bidirectional transport of Ca^2+^ into and out of mitochondria is a conserved biological process controlling multiple events, including metabolism, signaling, and cell fate. In the presence of membrane potential driving mitochondrial Ca^2+^ accumulation, transient changes of [Ca^2+^]_mt_ in response to cytosolic [Ca^2+^] variations are ensured by a molecular machinery for Ca^2+^ influx and efflux embedded in the inner mitochondrial membrane (IMM). While it is well established that influx relies on the Mitochondrial Calcium Uniporter (MCU), efflux was expected to be molecularly diversified, given the occurrence of functionally different exchange pathways with either Na^+^ or H^+1^. Accordingly, dedicated transporters ensure proper Ca^2+^ homeostasis and tightly regulated mitochondrial bioenergetics, but the process is not yet fully elucidated. We here demonstrate that TMEM65, a protein with an unknown biological function, is a fundamental component of the Ca^2+^ efflux machinery of mitochondria. As the MCU, TMEM65 has a broad tissue expression and localizes to the IMM. Its overexpression dramatically enhances Na^+^- and Li^+^-dependent mitochondrial Ca^2+^ extrusion, which is abrogated by the pharmacological inhibitor CGP-37157. Conversely, its downregulation chronically elevates resting mitochondrial Ca^2+^ levels and inhibits efficient Ca^2+^ efflux upon cellular activation, culminating in mitochondrial calcium overload and causing organelle dysfunction. Since TMEM65 has been associated with a severe human mitochondrial disease^2^, we deleted the TMEM65 homologues in *Caenorhabditis elegans* (*CeTMEM65*). While the two CeTMEM65 orthologs are dispensable for the survival at permissive growing conditions, their loss undermines embryonic developments when eggs are exposed to mild temperature-stress. In this regard, we find that *CeTMEM65 (null)* alleles cause necrotic lesions that are suppressed by inhibiting the mitochondrial calcium uniporter MCU-1. Overall, these results unambiguously assign a primary role in mitochondrial Ca^2+^ homeostasis to the orphan protein TMEM65. More importantly, our findings describe a novel molecular component that may be relevant in pathological settings in which excessive mitochondrial Ca^2+^ accumulation critically contribute to degenerative pathways.

---

Calcium (Ca^2+^) acts as a universal and ubiquitous intracellular messenger, exerting control over every facet of cellular pathophysiology^3^. Such a remarkable versatility relies on the one hand on a large array of dedicated Ca^2+^ binding and/or transporting proteins, and on the other hand on the strictly controlled compartmentalization of [Ca^2+^] changes. In such a sophisticated framework, mitochondria significantly shape intracellular Ca^2+^ dynamics and downstream Ca^2+^-dependent pathways. In healthy cells at rest, intramitochondrial [Ca^2+^] ([Ca^2+^]_mt_) matches cytoplasmic [Ca^2+^] ([Ca^2+^]_cyt_) at approximately 100 nM. Upon cellular stimulation, mitochondria undergo a rapid transient rise in [Ca^2+^]_mt_ that largely exceed bulk [Ca^2+^]_cyt_, up to 1 mM in a few cases^4,5^. This marked accumulation of Ca^2+^ is in part due to the highly negative organelle membrane potential (−180 mV), but also to the strategic positioning of mitochondria near Ca^2+^-releasing channels of the endoplasmic reticulum or the plasma membrane, where local micro/nano-domains of high [Ca^2+^] are generated^6^. Inside mitochondria, Ca^2+^ acts as a positive modulator of three dehydrogenases (PDH, IDH3 and OGDH), which stimulating oxidative metabolism and eventually ATP synthesis^7^. In pathological conditions, excessive Ca^2+^ accumulation within the organelle matrix (commonly referred to as Ca^2+^ overload) can potentially trigger permeability transition, ultimately leading to membrane permeabilization that may promote the release of pro-death factors^8–10^. Functioning as auxiliary cellular Ca^2+^ reservoirs, mitochondria contribute to buffering intracellular [Ca^2+^] transients, thereby influencing the spatio-temporal patterns of [Ca^2+^] oscillations, either within micro/nano-domains or on a bulk scale (e.g., throughout the entire cell)^11^. Consequently, impairment of mitochondrial Ca^2+^ homeostasis has been implicated in the pathogenesis of a variety of human diseases, ranging from neuromuscular diseases to brain injuries^12,13^.

From a biophysical perspective, [Ca^2+^]_mt_ is dynamically determined by the rates of Ca^2+^ influx versus efflux across the IMM. Pioneering investigations conducted in the 1960s and 1970s elucidated the mechanisms underlying this regulation: Ca^2+^ entry into mitochondria is facilitated by an electrogenic uniporter, while Ca^2+^ extrusion occurs through two distinct mechanisms - one dependent on sodium ions (i.e. mediated by a Na^+^/Ca^2+^ exchanger) and the other Na^+^-independent (i.e. mediated by a H^+^/Ca^2+^ exchanger)^14,15^. At the molecular level, mitochondrial Ca^2+^ influx is orchestrated by the MCU complex, whose precise composition varies across tissues and may encompass components such as MCU, MCUb, EMRE, MICU1, MICU2, and MICU3^16^. As to the efflux, the protein NCLX, encoded by the SLC8B1 gene, has been identified as the mediator of Na^+^-dependent Ca^2+^ extrusion^17^, whereas two distinct proteins, LETM1^18,19^ and TMBIM5^20,21^, have been proposed to function as mitochondrial H^+^/Ca^2+^ exchangers. Given the breakthrough discoveries of the last decade, it is not possible to rule out that other molecular components may contribute to mitochondrial Ca^2+^ transport. Here, we identify TMEM65 as a novel, unexpected, broadly conserved protein that enables Na^+^-dependent mitochondrial Ca^2+^ efflux. Genetic manipulation of TMEM65 expression has yielded striking effects on mitochondrial Ca^2+^ homeostasis in cultured cells. Specifically, overexpression of TMEM65 virtually eliminates [Ca^2+^]_mt_ transients, while its silencing prevents the decay of the organellar Ca^2+^ rises, resulting in markedly elevated resting [Ca^2+^]_mt_ levels. Further highlighting the prominent contribution of TMEM65 to organelle Ca^2+^ clearance, we show that genetic deletion of *TMEM65* homologues in *Caenorhabditis elegans* causes an overt necrotic phenotype that could be prevented by inhibiting mitochondrial Ca^2+^ influx, thereby pointing out that mitochondrial Ca^2+^ overload is a crucial determinant of cell demise. Since a human subject harboring a homozygous *TMEM65* loss-of-function mutation exhibits a neuromuscular phenotype reminiscent of mitochondrial disease^2^, it may be that MCU inhibition is a potential therapeutic strategy for conditions characterized by excessive mitochondrial Ca^2+^ accumulation.

## TMEM65 is a conserved protein of the IMM

We previously performed a bioinformatic analysis and identified candidate mitochondrial proteins of the inner membrane with a putative function in Ca^2+^ signaling^22^. Besides MCU, the list contains the recently described mitochondrial H^+^/Ca^2+^ exchanger TMBIM5 (also known as GHITM/MICS1). Thus, we sought to further inspect the list of presumed Ca^2+^ transporters using an evolutionary analysis to look for proteins with a conservation profile similar to MCU, and focused on a protein of unknown function named TMEM65 (Fig 1a). Among these candidates, TMEM65 is a 25 kDa protein with 3 transmembrane domains and a glycine zipper motif that are well conserved in kinetoplastids, green algae (but not in other plants) and metazoan (Fig 1b and Extended Data Fig 1a-b). Mammalian TMEM65 also shows an apparent poly-acidic tail near the C terminus. Although two works proposed TMEM65 as a protein positioned to cardiac sarcolemma^23,24^, multiple lines of evidence indicate that TMEM65 is a *bona fide* mitochondrial protein. In this regard, i) TMEM65 expression is not limited to the heart but rather is ubiquitous (highest in the cerebellum, lowest in the liver, Extended Data Fig 1c), ii) sequence analysis of TMEM65 reveals a conserved, prototypical mitochondrial targeting signal at the N-terminal (Fig 1b, predicted MPP-cleaved peptide indicated in cyan), and iii) TMEM65 is present in all high quality mitochondrial datasets based on robust proteomic evidence, including MitoCarta^25^, MitoCOP^26^ and MitoPhos^27^. To confirm these predictions and assess TMEM65 localization, we cloned, tagged (3xFlag) and expressed human TMEM65 in HeLa cells. Immunoblot analyses show a major band running with high specificity at 20 kDa, in line with the predicted molecular weight of the mature protein after processing by the Mitochondrial Processing Peptidase (MPP). The unprocessed 25 kDa precursor is also detectable upon overexpression (Fig 1c). Immunofluorescence analyses confirm co-localization with the mitochondrial marker HSP60 (Fig 1d). To test protein topology across the IMM, we performed Proteinase K (PK) protection assay in isolated mitoplasts. Digestion of intact (but not permeabilized) mitoplasts causes the loss of full-length TMEM65 and the appearance of a smaller fragment, indicating that a significant portion of TMEM65 resides within the matrix. Of note, the Flag peptide fused at the C-terminus of TMEM65 is lost after PK treatment (Fig 1e), thereby demonstrating that TMEM65 is inserted in the IMM as depicted in Fig 1b. This membrane orientation is also supported by other studies^28^, including an independent spatially-resolved proteomic screening that mapped Tyrosine 77 within the matrix (Y77, highlighted in grey in Fig 1b)^29^.

**Figure 1.**
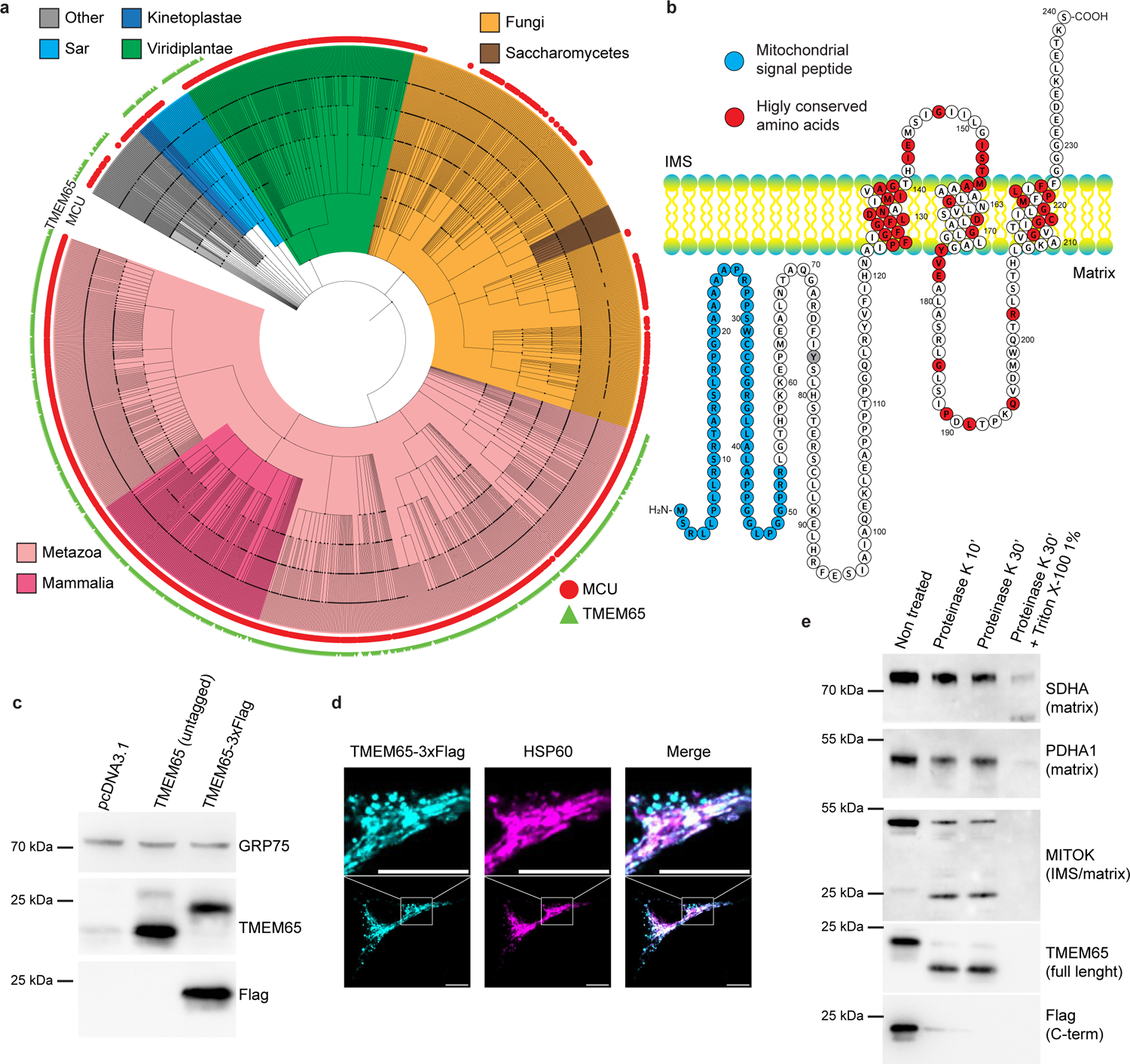
TMEM65 is a conserved protein of the IMM. (a) Cladogram showing the phylogenetic distribution of MCU (red) and TMEM65 (green) homologs across 1088 reference proteomes. (b) Schematic representation of TMEM65 membrane topology. (c) Western blot showing expression of TMEM65 and TME65-3xFlag in HeLa cells. (d) Immunolocalization of TMEM65-3xFlag (cyan) and the mitochondrial marker HSP60 (magenta) in HeLa cells, scale bar is 10 μm. (e) Proteinase K protection assay performed in mitoplasts isolated from TMEM65-3xFlag-expressing HeLa cells.

## TMEM65 expression modulates mitochondrial Ca^2+^ homeostasis

We next tested the role of TMEM65 on mitochondrial Ca^2+^ handling by monitoring [Ca^2+^]_mt_ in living intact HeLa cells using mitochondrial-targeted genetically-encoded Ca^2+^ indicators (GECI) based on the Ca^2+^-sensitive bioluminescent protein Aequorin^30^. Control and TMEM65-overexpressing cells were continuously perfused with modified Krebs-Ringer buffer and challenged with 100 μM histamine, an inositol 1,4,5-trisphosphate (IP_3_)-generating agonist that elicits Ca^2+^ release from the endoplasmic reticulum, leading to a transient increase of [Ca^2+^] in both cytoplasm and mitochondria. Interestingly, TMEM65 overexpression robustly inhibits (approximately 80%) the peak of [Ca^2+^]_mt_ upon cellular stimulation (Fig 2a). Importantly, this effect on mitochondrial Ca^2+^ handling is specific, since no major changes were detected in terms of cytosolic Ca^2+^ dynamics (Extended Data Fig 2a-b) and mitochondrial membrane potential (Extended Data Fig 2c), hence ruling out possible confounding factors. To further confirm the specificity of the effect and obtain additional insights into the structure-to-function relationship of TMEM65, we generated a set of mutants by introducing individual substitutions in the most conserved amino acids or by disrupting relevant motifs (i.e. glycine zipper and poly-acidic tail, Extended Data Fig 2d). All TMEM65 variants are expressed at a similar level, with the notable exception of the mutant lacking the transmembrane glycine zipper motif (GZ dead), that completely fail to express, probably due to an impairment in membrane packing (Extended Data Fig 2e)^31^. We evaluated histamine-induced [Ca^2+^]_mt_ peaks in cells expressing wild-type (wt) or mutant TMEM65. Our data (Extended Data Fig 2f-g) indicate that D132, D167, G170 and E178 are required for TMEM65 activity, while G140L and Q195E mutants are very similar to wt. The TMEM65 mutant lacking the poly-acidic tail still retain some effect, although to a lower extent when compared to wt. Surprisingly, the E144Q mutant shows even stronger effect than the wt. Overall, these data suggest a clear, specific involvement of TMEM65 in the regulation of mitochondrial Ca^2+^ fluxes.

**Figure 2.**
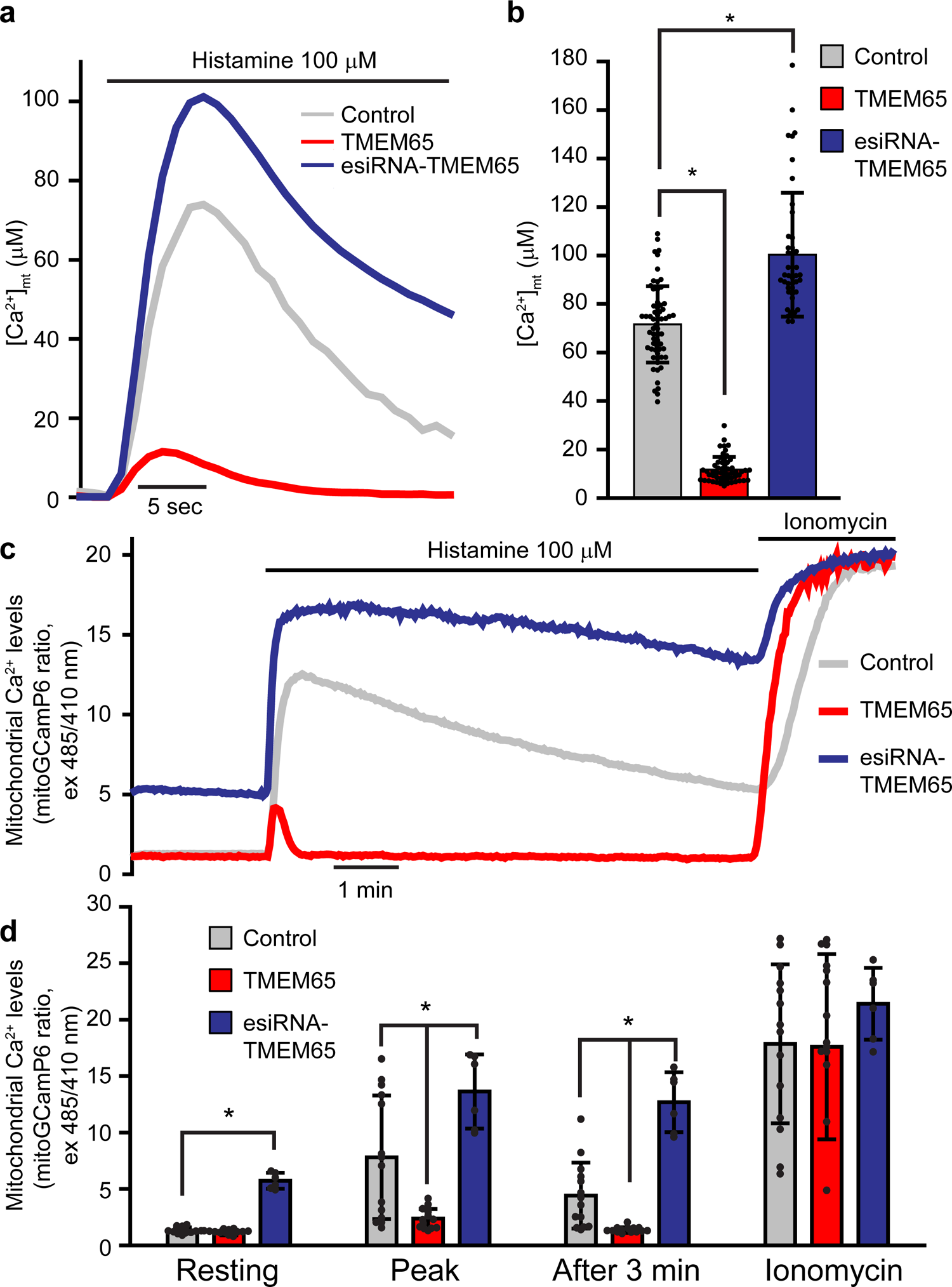
TMEM65 controls mitochondrial Ca^2+^ homeostasis. (a-b) [Ca^2+^]_mt_ measurements in HeLa cells expressing an aequorin-based probe. (a) Representative [Ca^2+^]_mt_ traces of control, TMEM65 overexpressing and silenced cells. (b) Histograms (mean ± s.d.) of histamine-induced [Ca^2+^]_mt_ peak values. (c-d) [Ca^2+^]_mt_ measurements in HeLa cells expressing the mitoGCaMP6 probe. (c) Representative traces of control, TMEM65 overexpressing and silenced cells. (d) Histograms (mean ± s.d.) of mitochondrial Ca^2+^ levels at the indicate time points. * indicates *p* < 0.001 using one-way ANOVA with Holm-Sidak correction. Individual data points, descriptive statistics and exact *p* values are included in Source Data Table.

We next evaluated the effect of TMEM65 downregulation on organelle Ca^2+^ dynamics by testing different synthetic unmodified siRNAs. When agonist-induced peak [Ca^2+^]_mt_ values were compared, we obtained inconsistent results among siRNA sequences (Extended Data Fig 3a-b) despite comparable efficiency in reducing TMEM65 protein level (Extended Data Fig 3e), an outcome likely due to off-target effects. To minimize non-specificity, we used endoribonuclease-prepared siRNAs (esiRNA)^32^. This approach reveals increased [Ca^2+^]_mt_ peak upon specific downregulation of TMEM65 (Fig 2a-b). Most importantly, we also observed that, in addition to the effect on maximal [Ca^2+^]_mt_ values, TMEM65 silencing alters the kinetic profile of [Ca^2+^]_mt_ transients. In particular, a delayed Ca^2+^ clearance after the peak is noticeable, an indication of impaired mitochondrial Ca^2+^ extrusion. To better appreciate this effect, we re-analyzed all Ca^2+^ traces by normalizing [Ca^2+^]_mt_ on peak [Ca^2+^] (Extended Data Fig 3c). The slope of the resulting trace is thus proportional to mitochondrial Ca^2+^ efflux rate (Extended Data Fig 3d). This analysis clearly reveals that TMEM65 overexpression enhances mitochondrial Ca^2+^ extrusion (notwithstanding lower [Ca^2+^]_mt_ peak), while its silencing elicits the opposite effect. Remarkably, all tested siRNAs consistently decreases mitochondrial Ca^2+^ efflux, irrespective of their wavering effect on peak [Ca^2+^]_mt_ (Extended Data Fig 3a-d). Altogether, these data suggest an involvement of TMEM65 in mitochondrial Ca^2+^ efflux rather than in influx. To further substantiate our data, we evaluated mitochondrial Ca^2+^ dynamics using a GFP-based GECI, the mitoGCaMP6 probe, that also allows the more precise quantification of baseline [Ca^2+^]_mt_ and extrusion kinetics^33^. Compared to controls, both overexpression and downregulation of TMEM65 alter mitochondrial Ca^2+^ dynamics, further supporting that TMEM65 is a key regulator of mitochondrial Ca^2+^ efflux. Specifically, TMEM65 overexpression causes a negligible decrease of resting Ca^2+^, a histamine induced upstroke similar to control, at least in the initial phase, but most evidently, a steep return to baseline after the peak (Fig 2c). Conversely, TMEM65-silenced cells show chronically elevated resting matrix Ca^2+^ levels, and histamine-induced [Ca^2+^] elevations that last over several minutes (Fig 2c-d). Altogether, our detailed analyses of mitochondrial Ca^2+^ dynamics uncover a prominent role of TMEM65 in controlling mitochondrial Ca^2+^ efflux.

## TMEM65 mediates Na^+^-dependent mitochondrial Ca^2+^ efflux

We next explore the mechanism underlying TMEM65 effect on mitochondrial Ca^2+^. It is known that experiments carried out in intact cells have the benefit of examining mitochondria in their native environment. However, these experimental settings suffer of two main drawbacks. Firstly, mitochondrial Ca^2+^ influx and efflux constantly coexist, thus preventing the genuine segregation of the two phenomena. Secondly, efflux relies on two concurrent but separate mechanisms, one Na^+^-dependent and the other Na^+^-independent. To overcome these limitations, we performed [Ca^2+^]_mt_ measurements in digitonin-permeabilized cells, where mitochondria can be exposed to buffers with defined ionic composition. In this experimental setup, mitochondrial Ca^2+^ uptake is initiated by perfusing mitochondria of plasma-membrane permeabilized cells with an intracellular-like buffer containing fixed [Ca^2+^], until a plateau [Ca^2+^]_mt_ is reached (i.e. when influx rate equals efflux rate). Then, Ruthenium Red (RuR), the best-known MCU inhibitor, is added to block the influx, thus causing a progressive decrease of [Ca^2+^]_mt_. The rate of this decrease represents a reliable measurement of mitochondrial Ca^2+^ efflux. Perfusion with Na^+^-free buffer allows the evaluation of organelle Na^+^-independent efflux, while incorporation of 10 mM NaCl in the RuR-containing buffer enhances mitochondrial Ca^2+^ extrusion, thereby enabling the quantification of Na^+^-dependent efflux. Li^+^ can also be used as Na^+^ surrogate, although less effective^34^. Using this approach, we found that both TMEM65 overexpression and downregulation show no effects on neither mitochondrial Ca^2+^ influx nor Na^+^-independent efflux. Rather, TMEM65 overexpression greatly enhances Na^+^- and Li^+^-dependent mitochondrial Ca^2+^ efflux. Accordingly, its silencing blunts the Na^+^-dependent component of organellar Ca^2+^ extrusion (Fig 3a-d, Extended Data Fig 4a-b). To further support this finding, we took advantage of the specific pharmacological inhibitor of mitochondria Na^+^/Ca^2+^ exchange, the benzothiazepine CGP-37157^35^. This compound causes an increase in agonist-induced [Ca^2+^]_mt_ peaks in control cells due to the inhibition of organelle Ca^2+^efflux^36^. Strikingly, CGP-37157 is able to abrogate the effect elicited by TMEM65 overexpression, restoring normal mitochondrial Ca^2+^ dynamics (Fig 3 e-g). Overall, these results indicate that i) mitochondrial Ca^2+^ efflux strongly counteracts both in resting conditions and in all phases of agonist-evoked responses the action of MCU and ii) TMEM65 is an essential component and/or the strongest contributor to the process of mitochondrial Ca^2+^ efflux.

**Figure 3.**
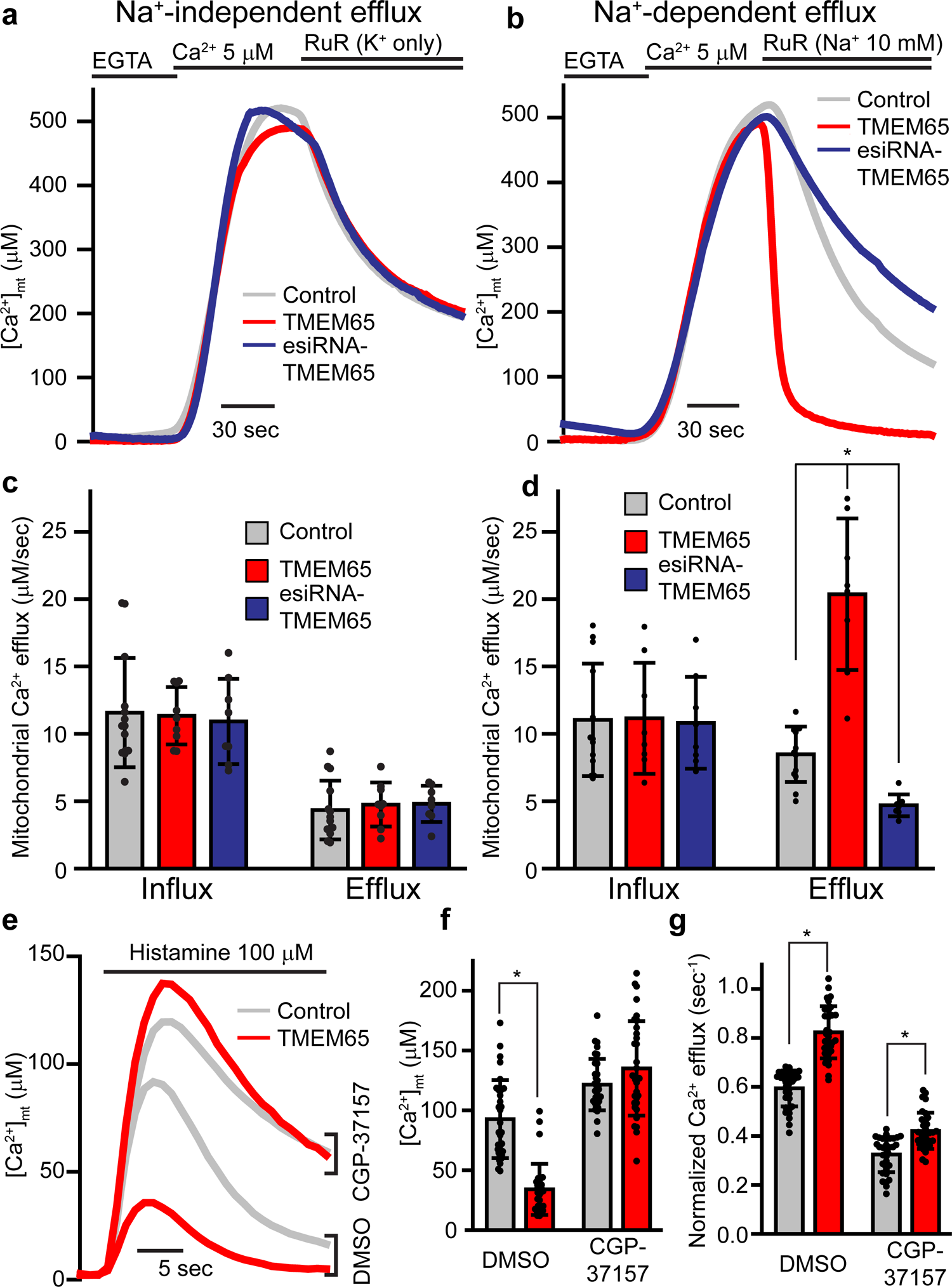
TMEM65 mediates Na^+^-dependent mitochondrial Ca^2+^ efflux. (a-d) [Ca^2+^]_mt_ measurements in digitonin-permeabilized HeLa cells. (a-b) Representative [Ca^2+^]_mt_ traces of control, TMEM65 overexpressing and silenced cells in the absence (a) or presence (b) of 10 mM Na^+^. (c-d) Histograms (mean ± s.d.) of maximal Na^+^-independent (c) and Na^+^-dependent (d) mitochondrial Ca^2+^ influx and efflux rates. * indicates *p* < 0.001 using one-way ANOVA with Holm-Sidak correction. (e-g) [Ca^2+^]_mt_ measurements in control and TMEM65 overexpressing intact HeLa cells treated with either vehicle (DMSO) or 20 μM CGP-37157. (e) Representative [Ca^2+^]_mt_ traces. (f) Histograms (mean ± s.d.) of histamine-induced [Ca^2+^]_mt_ peak values. (g) Histograms (mean ± s.d.) of normalized mitochondrial Ca^2+^ efflux. * indicates *p* < 0.001 using two-way ANOVA with Holm-Sidak correction. Individual data points, descriptive statistics and exact *p* values are included in Source Data Table.

## TMEM65 deficiency causes necrotic lesions *in vivo* in *C. elegans*

To better understand the physiological relevance of TMEM65, we set out a series of experiments using the nematode *Caenorhabditis elegans* as an *in vivo* model organism. Protein alignment tools predict that *C33H5.19/tag-321* and *C03B8.3* genes encode for two TMEM65 orthologs (here referred to also as CeTMEM65-1 and CeTMEM65-2, respectively) with 52% and 35% similarities respectively (Extended Data Fig 5a-c). First, we tested the expression and functionality of CeTMEM65 in mammalian cells. Western blot shows that CeTMEM65-1 is efficiently expressed also in HeLa cells, while CeTMEM65-2 fails to be detected, probably due to instability of the protein at 37°C (Extended Data Fig 5d). In terms of activity, the effect of CeTMEM65-1 on mitochondrial Ca^2+^ handling is comparable to its human counterpart, indicating that their biochemical function is maintained through evolution (Extended Data Fig 5e). Then, we generated *CeTMEM65* mutants by using a CRISPR/Cas9 approach previously validated in some of our studies^37–40^. We obtained *tag-321(bon103)* and *C03B8.3(bon104)*, two alleles that are predicted to be *(null)* because both genetic lesions deleted portions of the promoters as well as all exons encoding for the corresponding proteins (Extended Data Fig 6a). While *tag-321(bon103)* and *C03B8.3(bon104)* single mutants do not display obvious difference compared to wt (N2) nematodes, *C03B8.3;tag-321* double *(null)* mutants had a slight developmental delay that was accompanied by a mild, though significant reduction of hatched larvae at 20 °C (Fig 4a). Preliminary evidence showed no major survival difference between wt and *CeTMEM65* single and double mutant nematodes (Extended Data Fig 6b and Supplementary Table 1). To better characterize our model, we performed label free quantitative liquid chromatography-tandem mass spectrometry (LC-MS/MS) using samples from wt *and CeTMEM65 (null)* young adults grown at 20 °C. Compared to control, we found 147 differentially expressed proteins (Fig 4b), many of which are involved in lipid metabolism (9 DEP) and autophagy/endocytosis (3 DEP in autophagosome assembly, 4 DEP in endocytosis) (Extended Data Fig 6c). To test the influence of CeTMEM65 on lipid content, we generated a strain expressing the DHS-3::GFP fusion reporter. DHS-3 is a short-chain dehydrogenase which has been found to be highly abundant in lipid droplets in *C. elegans* intestine^41,42^. We then performed confocal imaging along with western blot analysis and found an increased amount of GFP-positive lipid droplets in *CeTMEM65* double mutants, suggesting that loss of CeTMEM65 might influence lipid biosynthesis or catabolism (Figure 4c and Extended Data Fig 6d). Since lipid metabolism may contribute to *C. elegans* survival and adaptation when animals are shifted to low or high temperatures^43,44^, we performed a series of lifespan assays at 25 °C, a temperature known to influence *C. elegans* development, survival and stress response^45^. Both *C03B8.3 (null)* and *tag-321 (null)* single mutants had a median lifespan that was comparable to wt nematodes, whereas *C03B8.3;tag-321* double mutant nematodes lived slightly less (Extended Data Fig 6e and Supplementary Table 1). When we had a closer look at plates of animals grown at 25 °C, we noticed that most of the *CeTMEM65 (null)* eggs did not hatch and, more importantly, they displayed large vacuoles (Fig 4d), resembling necrotic structures, very similar to those necrotic lesions observed in neurons expressing degenerin channels^46,47^. We sought to further substantiate our initial observation by quantifying the number of animals that could hatch from eggs laid by nematodes grown for a single generation at 25 °C (Fig 4e). Intriguingly, we found that only a handful of *C03B8.3;tag-321* double mutants could properly develop and reach adulthood (Fig 4f). Of note, *C03B8.3 (null)* mutants had a hatching rate similar to wt, whereas *tag-321 (null)* animals showed a modest, although significant increased number of larvae that could survive at 25 °C (Fig 4f). Together, this severe degenerative phenotype is consistent with the proposed molecular function of TMEM65. Indeed, mitochondrial Ca^2+^ extrusion in *CeTMEM65 (null)* cells should be dramatically reduced, thereby leading to organellar Ca^2+^ overload accompanied by extensive cell death. Because of that, we reasoned that inhibition of mitochondrial Ca^2+^ influx (e.g. by deletion of nematode *mcu-1* gene) in *CeTMEM65 (null)* background should prevent mitochondrial Ca^2+^ overload and restore normal hatching at 25 °C. We first attempted to generate *C03B8.3;mcu-1;tag-321* triple mutants using the already existing alleles. However, since *mcu-1* and *tag-321* genes are both on chromosome IV at a very close distance (∼2Mb), we could not introduce the *mcu-1(ju1154)* allele^48,49^ into the existing *C03B8.3(null);tag-321(null)* mutants. To overcome this problem, we performed another gene editing in *mcu-1(ju1154)* animals and generated an equivalent lesion within *tag-321* gene. DNA sequencing of the obtained mutation confirmed that the new *bon105* allele disrupted the promoter and coding region of *tag-321* gene as the previous *bon103* allele (Extended Data Fig 6f). We backcrossed the new strain, generated *C03B8.3(bon104);mcu-1(ju1154);tag-321(bon105)* triple mutants and carried out another hatching assay. Remarkably, we found that *mcu-1* loss-of-function *(lof)* almost suppressed the degeneration of *C03B8.3;tag-321* double mutant eggs at 25 °C (Fig 4g-h), indicating an epistatic relationship between MCU and TMEM65 and supporting the prominent role of TMEM65 as a key regulator of mitochondrial Ca^2+^ efflux. Together, our *in vivo* experimental evidence demonstrates that TMEM65 deficiency may facilitate mitochondrial Ca^2+^ overload, when cells are exposed to environmental challenges (e.g., restrictive temperatures) that may deplete resources (e.g., lipids) and induce degenerative processes (necrosis).

**Figure 4.**
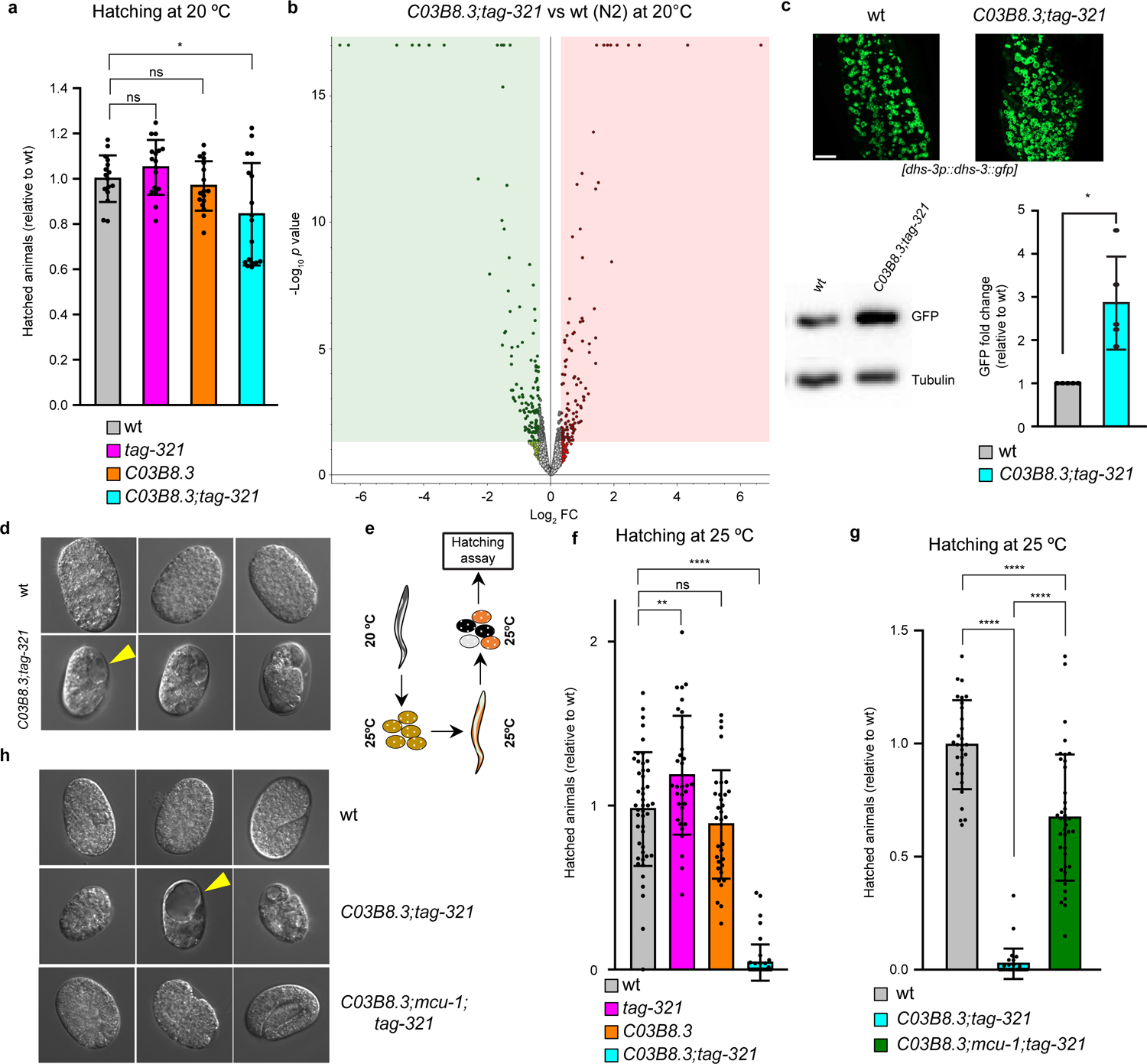
Loss of CeTMEM65 is detrimental for embryos development. (a) Quantification of hatched animals when eggs were seeded at 20 °C (n=2, with at least 8 adults used for each genotype. Ordinary one-way ANOVA followed by Dunnett’s multiple comparisons test. * indicates *p* < 0.05, ns= not significant). (b) Volcano plot of differentially regulated proteins using samples from *C03B8.3(null);tag-321(null)* mutants and wt nematodes grown at 20 °C. Significant protein expression is based on *p* value ≤ 0.05 and fold change ± 1.25 thresholds. (c) Upper panel shows representative confocal images of lipid droplets in wt and *C03B8.3(null);tag-321(null)* nematodes expressing DHS-3::GFP. Scale bar = 10μm. Lower panels show representative Western blot and densitometric analyses of wt and *C03B8.3(null);tag-321(null)* mutants adult nematodes carrying the DHS-3::GFP fusion reporter. Worms were grown at 20 °C until day 4. Mann Whitney test, * indicates *p* < 0.05. (d) Representative DIC images of eggs extracted from gravid animals grown at 25 °C. (e) Schematic representation of the hatching assay employed in this study. (f) Quantification of animals that hatched from eggs at 25 °C using the protocol describe in (e). The graph represents three independent biological replicates, in which at least 8 adult animals were used to determine the surviving progeny. Ordinary one-way ANOVA was followed by Turkey’s multiple comparisons test. ***p<0.001, **p<0.01, ns= not significant. (g) Hatching assay of eggs at 25 °C. The graph represents three independent biological replicates in which at least 8 adult animals were used to determine the surviving progeny. Ordinary one-way ANOVA was followed by Dunnett’s multiple comparisons test (****p<0.0001). (h) Representative DIC images of eggs extracted from gravid animals grown at 25 °C. Individual data points, descriptive statistics and exact *p* values are included in Source Data Table.

In conclusion, we herein unveil the previously undescribed function of TMEM65, by demonstrating its involvement in controlling mitochondrial Ca^2+^ fluxes. Our data suggest that TMEM65 is on the one hand sufficient to greatly enhance Na^+^-dependent mitochondrial Ca^2+^ extrusion, as demonstrated by overexpression experiments. On the other hand, TMEM65 is also necessary to prevent mitochondrial Ca^2+^ overload, as demonstrated by both *in vitro* experiments with siRNAs and *in vivo* using *C. elegans*. Our results also shed light on the pathogenic mechanism of a case report showing that TMEM65 deficiency causes mitochondrial dysfunction and leads to a severe neuromuscular syndrome in humans^2^. Based on our *in vivo* experimental paradigm, it seems that MCU inhibition may be a strategy to ameliorate pathogenic processes associated with TMEM65 deficiency. Considering that excessive mitochondrial Ca^2+^ has been implicated in several diseases, the present work sets the stage for the development of therapeutic approaches that focus on correcting mitochondrial Ca^2+^ overload.

## Acknowledgments

The authors are grateful to Luca Barbato (UniPD) for the help with mutagenesis, Ms. Christiane Bartling-Kirsch (DZNE) and Mr Amal John Mathew (DZNE) for their technical assistance. Moreover, we would like to thank Dr. Christopher E. Hopkins and Dr. Ben Jussila (InVivo Biosystems) for their technical support with CRISPR/Cas9 in C. elegans. This work was supported by grants from the University of Padova (STARS@UNIPD), the Italian Ministry of University and Research (PRIN 2017YF9FBS_002, 2020RRJP5L_002 and PRIN 2020R2BP2E_002), European Union (Next Generation EU CN00000041), the Italian Association for Cancer Research (AIRC 5 per mille 2019 - ID 22759), CARIPARO (ID 59583), CARIPLO and Telethon (GJC21054) foundations. DB is a member of the DFG Cluster of Excellence ImmunoSensation funded by the Deutsche Forschungsgemeinschaft (DFG, German Research Foundation) under Germany’s Excellence Strategy – EXC2151 – 390873048.

## Author information

All authors declare no competing interests.

## Author contributions

Massimo Vetralla: Methodology, Validation, Investigation, Formal analysis.

Lena Wischhof: Methodology, Validation, Formal analysis, Investigation, Writing-Original Draft, Visualization.

Vanessa Cadenelli: Validation, Investigation, Formal analysis.

Enzo Scifo: Methodology, Validation, Formal analysis, Investigation, Writing-Original Draft, Visualization.

Dan Ehninger: Resources, Funding acquisition.

Rosario Rizzuto: Conceptualization, Writing-Original Draft, Visualization, Supervision, Project administration, Funding acquisition.

Daniele Bano: Conceptualization, Methodology, Validation, Formal analysis, Investigation, Writing-Original Draft, Visualization, Supervision, Project administration, Funding acquisition. Diego De Stefani: Conceptualization, Methodology, Validation, Formal analysis, Investigation, Writing-Original Draft, Visualization, Supervision, Project administration, Funding acquisition.

## Extended Data Figure legends

**Extended Data Figure 1.**
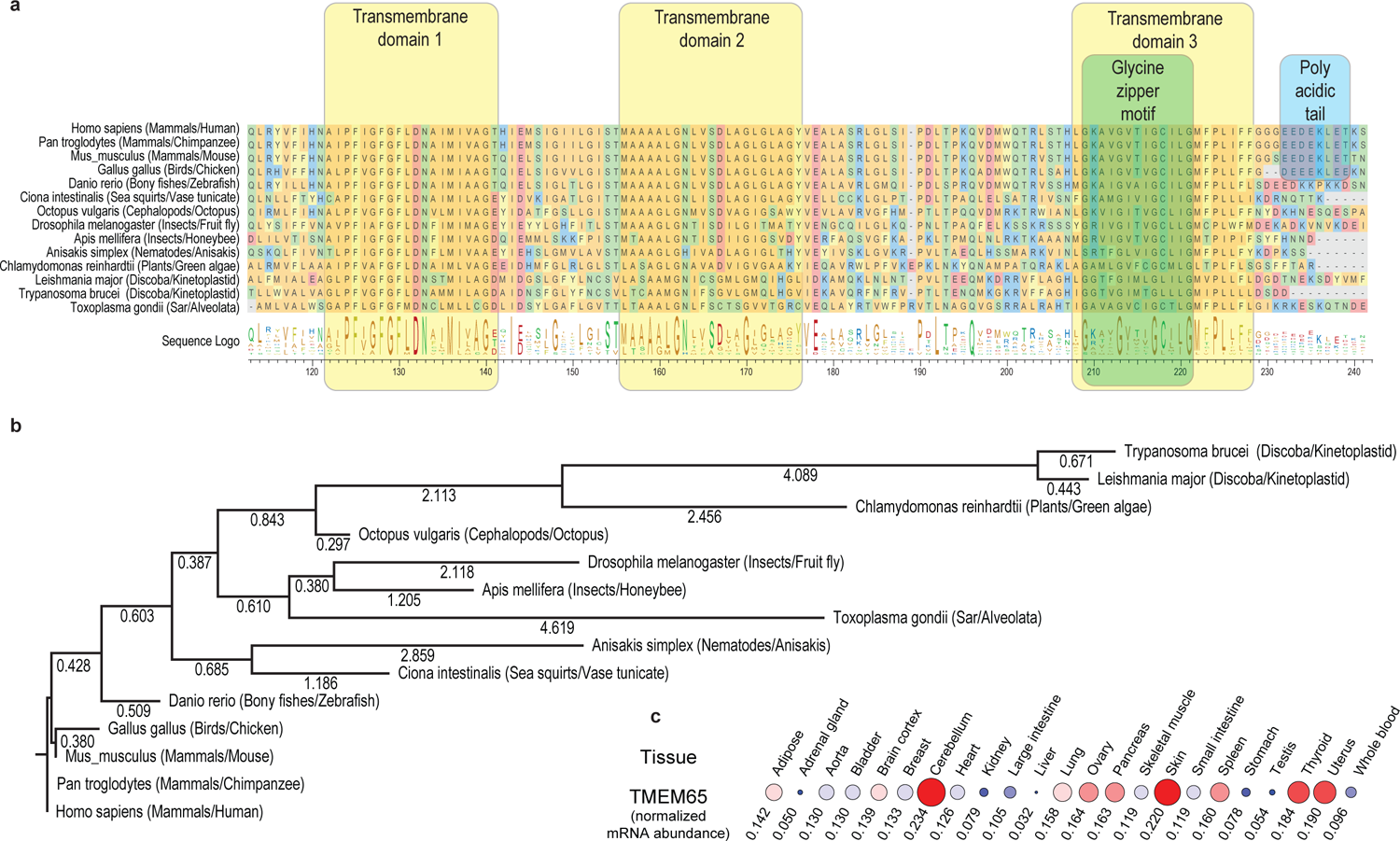
(a) Multiple sequence alignment of TMEM65 homologs from the indicated species. (b) Maximum Likelihood phylogenetic trees of selected TMEM65 homologs. (c) TMEM65 mRNA expression across the indicated human tissues. Data are normalized on the average abundance of nuclear-encoded transcripts of mitochondrial proteins.

**Extended Data Figure 2.**
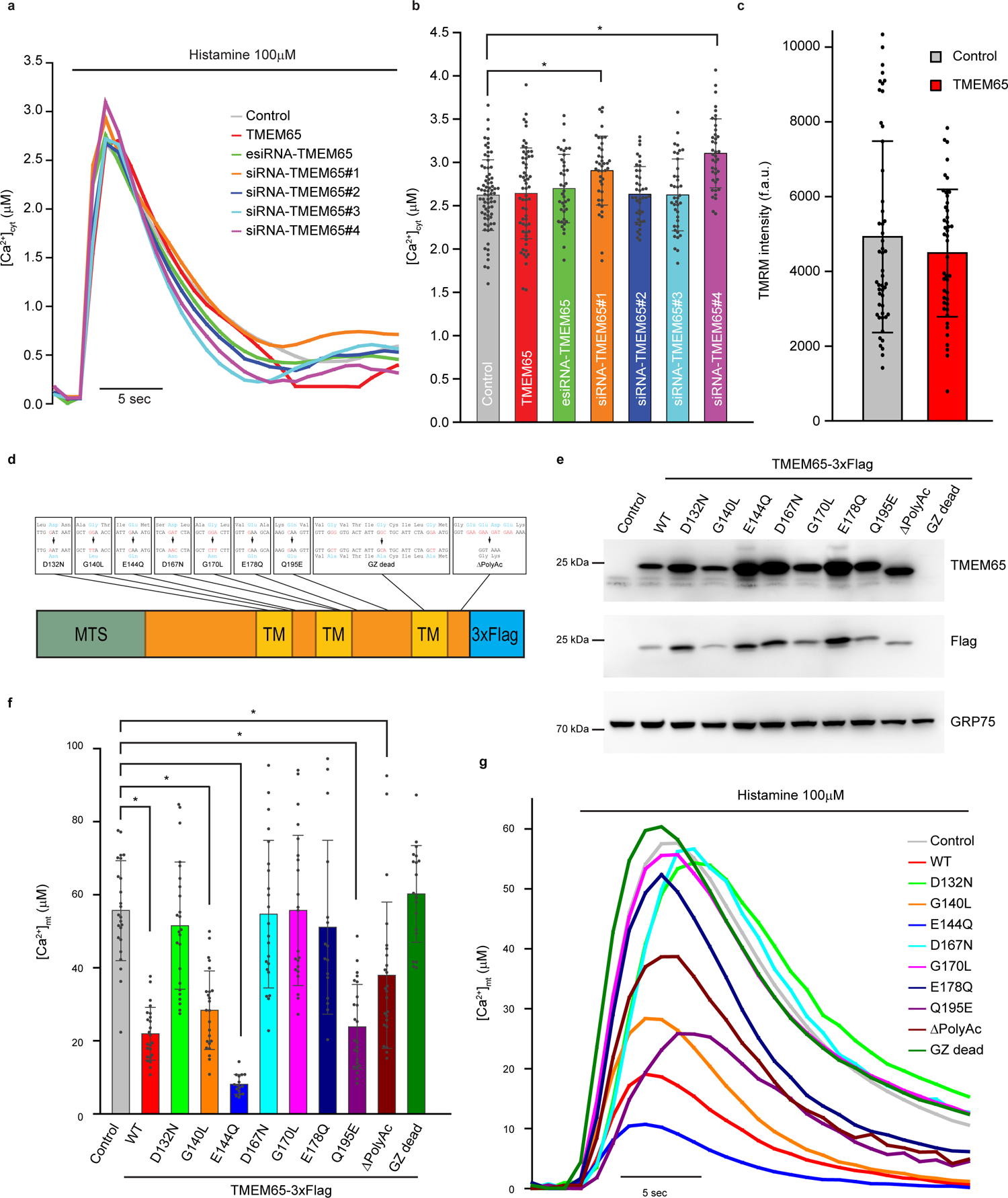
(a-b) [Ca^2+^]_cyt_ measurements in HeLa cells expressing an aequorin-based probe. (a) Representative [Ca^2+^]_cyt_ traces cells transfected with indicated constructs. (b) Histograms (mean ± s.d.) of histamine-induced [Ca^2+^]_mt_ peak values. (c) Basal TMRM intensities of control and TMEM65-overexpressing HeLa cells. (d) Schematic representation of TMEM65 mutants tested in this study. (e) Western blot of proteins extracted from HeLa cells transfected with wild type or TMEM65 variants. (f-g) [Ca^2+^]_mt_ measurements in HeLa cells expressing an aequorin-based probe. (f) Representative [Ca^2+^]_mt_ traces of cells expressing the indicated TMEM65 variants. (g) Histograms (mean ± s.d.) of histamine-induced [Ca^2+^]_mt_ peak values. * indicates *p* < 0.001 using one-way ANOVA with Holm-Sidak correction. Individual data points, descriptive statistics and exact *p* values are included in Source Data Table.

**Extended Data Figure 3.**
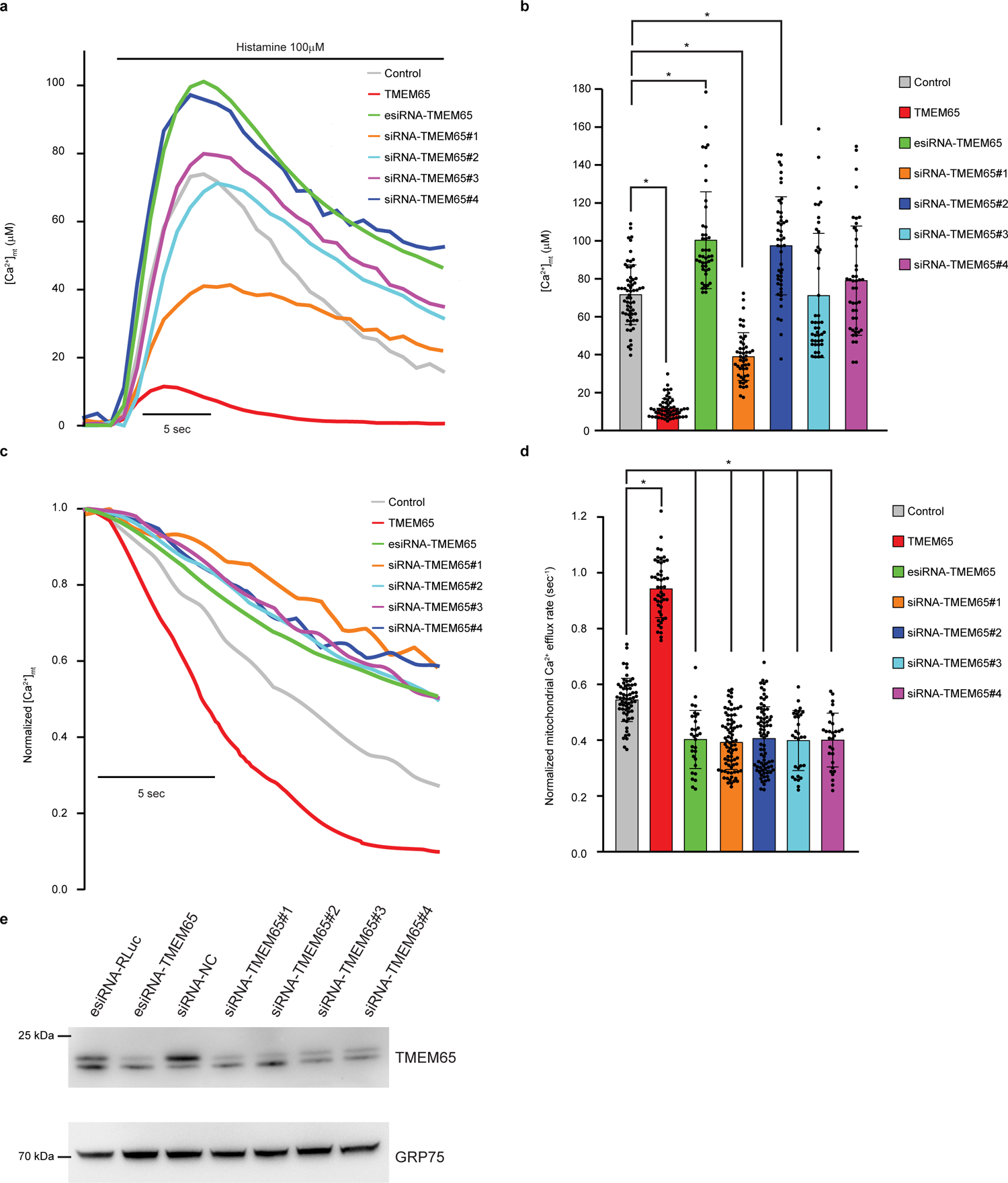
(a-d) [Ca^2+^]_mt_ measurements in HeLa cells expressing an aequorin-based probe. (a) Representative [Ca^2+^]_mt_ traces of cells transfected with the indicated construct. (b) Histograms (mean ± s.d.) of histamine-induced [Ca^2+^]_mt_ peak values. (c) Representative traces of normalized mitochondrial Ca^2+^ efflux. (d) Histograms (mean ± s.d.) of normalized mitochondrial Ca^2+^ extrusion rate. (e) Western blot of proteins extracted from HeLa cells transfected with the indicated constructs. * indicates *p* < 0.001 using one-way ANOVA with Holm-Sidak correction. Individual data points, descriptive statistics and exact *p* values are included in Source Data Table.

**Extended Data Figure 4.**
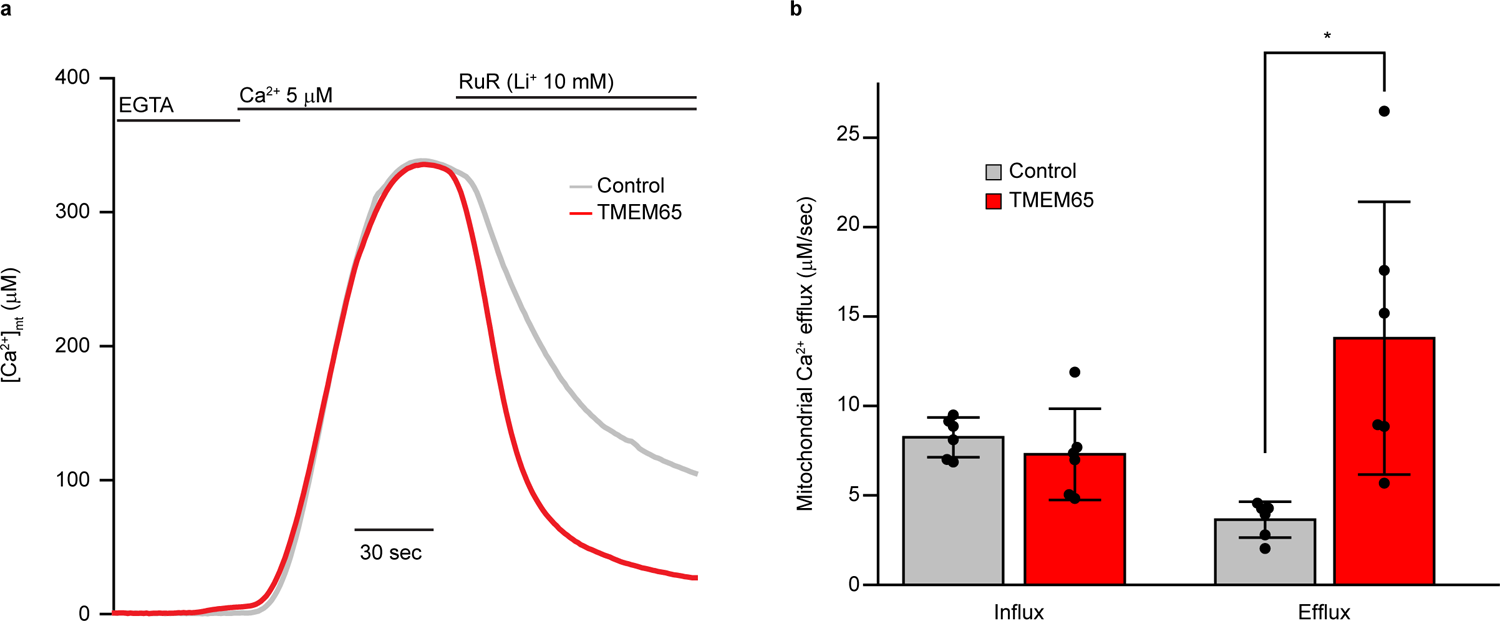
(a-b) [Ca^2+^]_mt_ measurements in digitonin-permeabilized HeLa cells. (a) Representative [Ca^2+^]_mt_ traces of control and TMEM65 overexpressing cells in the presence of 10 mM Li^+^. (b) Histograms (mean ± s.d.) of maximal Li^+^-dependent mitochondrial Ca^2+^ influx and efflux rates. Individual data points, descriptive statistics and exact *p* values are included in Source Data Table.

**Extended Data Figure 5.**
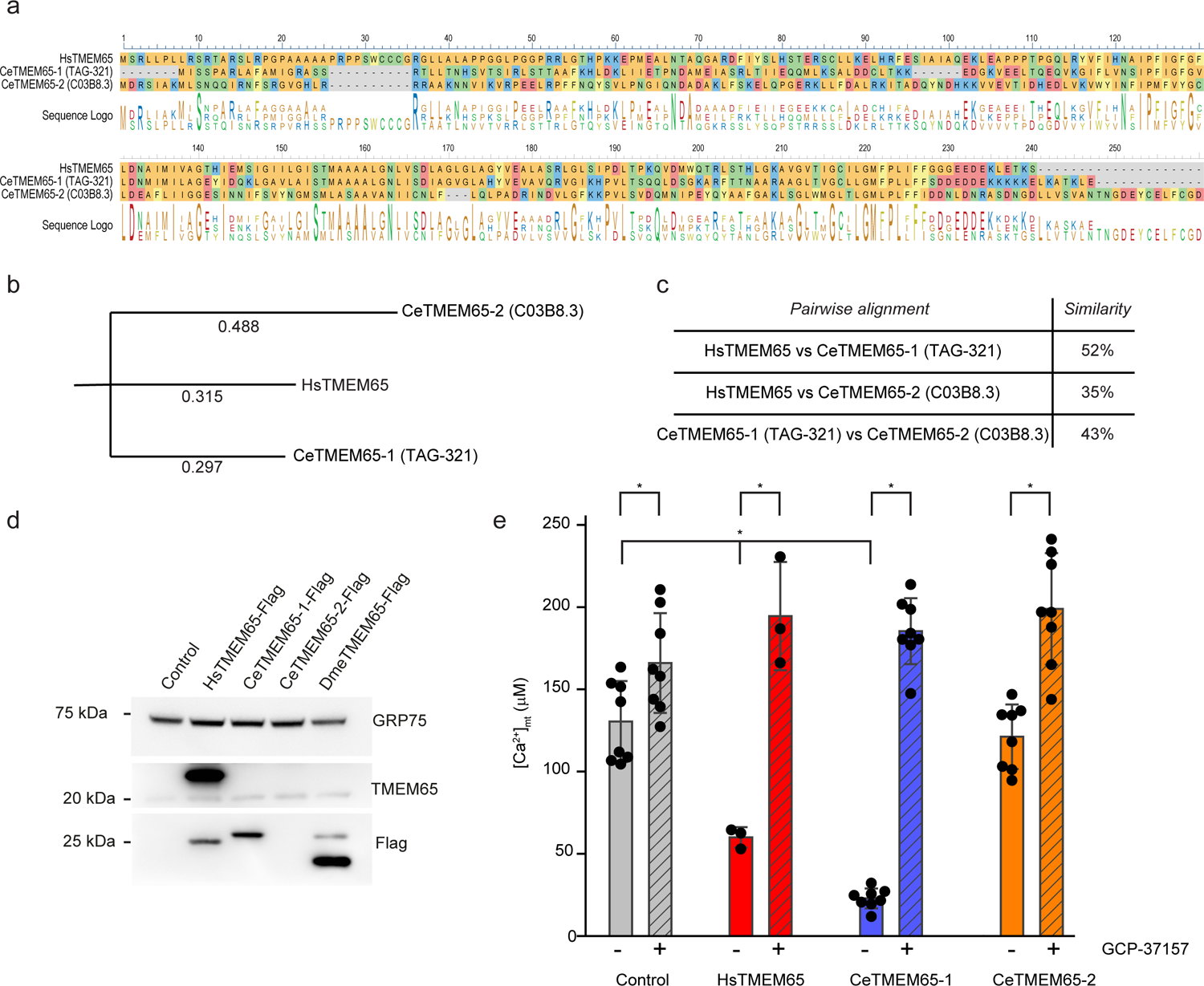
(a) Multiple sequence alignment of TMEM65 from H. sapiens and C. elegans (b) Maximum likelihood phylogenetic trees of selected TMEM65 homologs. (c) Sequence similarities between human and nematode TMEM65 based on pairwise alignments. (d) Western blot of proteins extracted from HeLa cells transfected with *H. sapiens* and *C. elegans* TMEM65. (e) Histograms (mean ± s.d.) of histamine-induced [Ca^2+^]_mt_ peak values recorded in HeLa cells transfected with *H. sapiens* and *C. elegans* TMEM65. Where indicated, cells were treated with 20 μM CGP-37157. Individual data points, descriptive statistics and exact *p* values are included in Source Data Table.

**Extended Data Figure 6.**
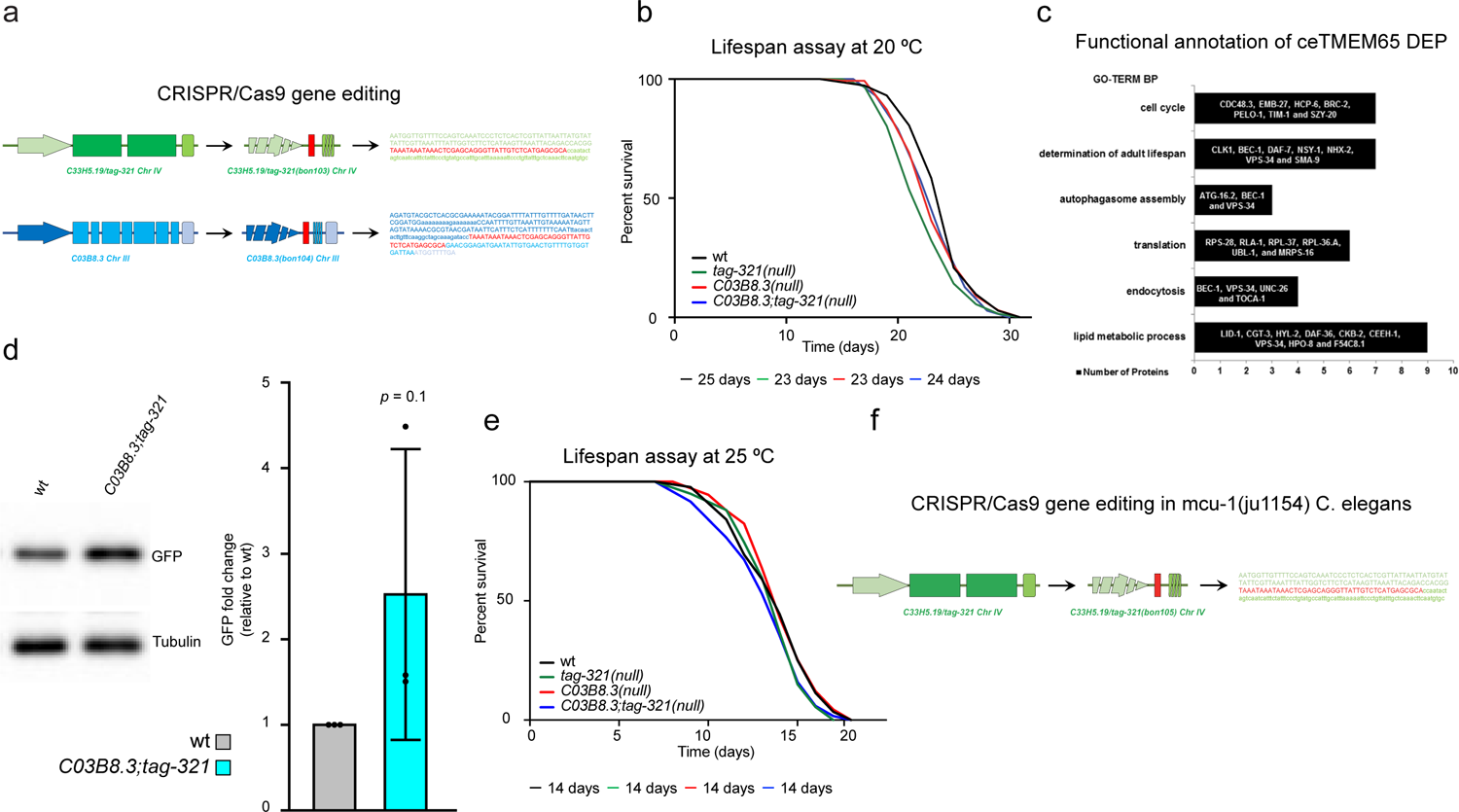
(a) Schematic representation of CRISPR/Cas9 gene editing on *tag-321* and *C03B8.3* genes. The DNA sequences of the two alleles are reported. (b) Lifespan assay of animals grown at 20 °C. (c) Functional annotation of differentially expressed proteins (DEP) in CeTMEM65 mutant nematodes compared to wt. (d) Representative Western blot and densitometric analyses of wt and *C03B8.3(null);tag-321(null)* mutants adult nematodes carrying the DHS-3::GFP fusion reporter. Worms were grown at 25 °C until day 4. Mann Whitney test, exact *p* is indicated. (e) Representative survival curves of wt, *tag-321*, *C03B8.3* and *C03B8.3;tag-321* mutant nematodes at 25 °C. Descriptive statistics, individual assays and exact *p* values are included in Supplementary table 1. (f) Schematic representation of CRISPR/Cas9 gene editing of *tag-321* allele in *micu-1(ju1154)* mutant nematodes. DNA sequence of the *tag-321(bon105)* allele is reported.

Supplementary Information is available for this paper

## Methods

### Chemicals, cell culture and transfection

All chemicals were purchased from Sigma-Aldrich, unless otherwise specified. All the experiments were performed in HeLa cells (ATCC Number: CCL-2) cultured in Dulbecco’s modified Eagle’s medium (DMEM) (Gibco #41966052, Thermo Fisher Scientific), supplemented with 10% fetal bovine serum (FBS) (Thermo Fisher Scientific), containing penicillin (100 U/ml) and streptomycin (100μg/ml) (Euroclone). When needed, cells were seeded on either glass coverslips (onto 13- or 24-mm in diameter) or 96-well plates and allowed grow to 50-80% confluence before transfection. Transfection of siRNA was performed using RNAiMAX (Thermo Fisher Scientific) according to manufacturer’s instructions. Transfection of plasmid DNA was performed using polyethylenimine (PEI MAX, Polysciences).

### Plasmids and siRNAs

Plasmids encoding C-term 3xFlag tagged human TMEM65 (NM_194291.2), *C. elegans* TAG-321 (NM_068883.5), and *C. elegans* C03B8.3 (NM_066278.5) were purchased from VectorBuilder. Untagged human TMEM65 was amplified using the following primer: fw 5’-ATTAGCTAGCACCACCATGTCCCGGCTG-3’, rv 5’-ATTAGAATTCTTAACTTTTCGTTTCCAGTTTTTCATCTTCTTC-3’. The PCR fragment was cloned into pcDNA3.1 (Thermo Fisher Scientific) using NheI and BamHI sites.

Mutagenesis was performed using In-Fusion cloning (Takara) using the following primer pairs:

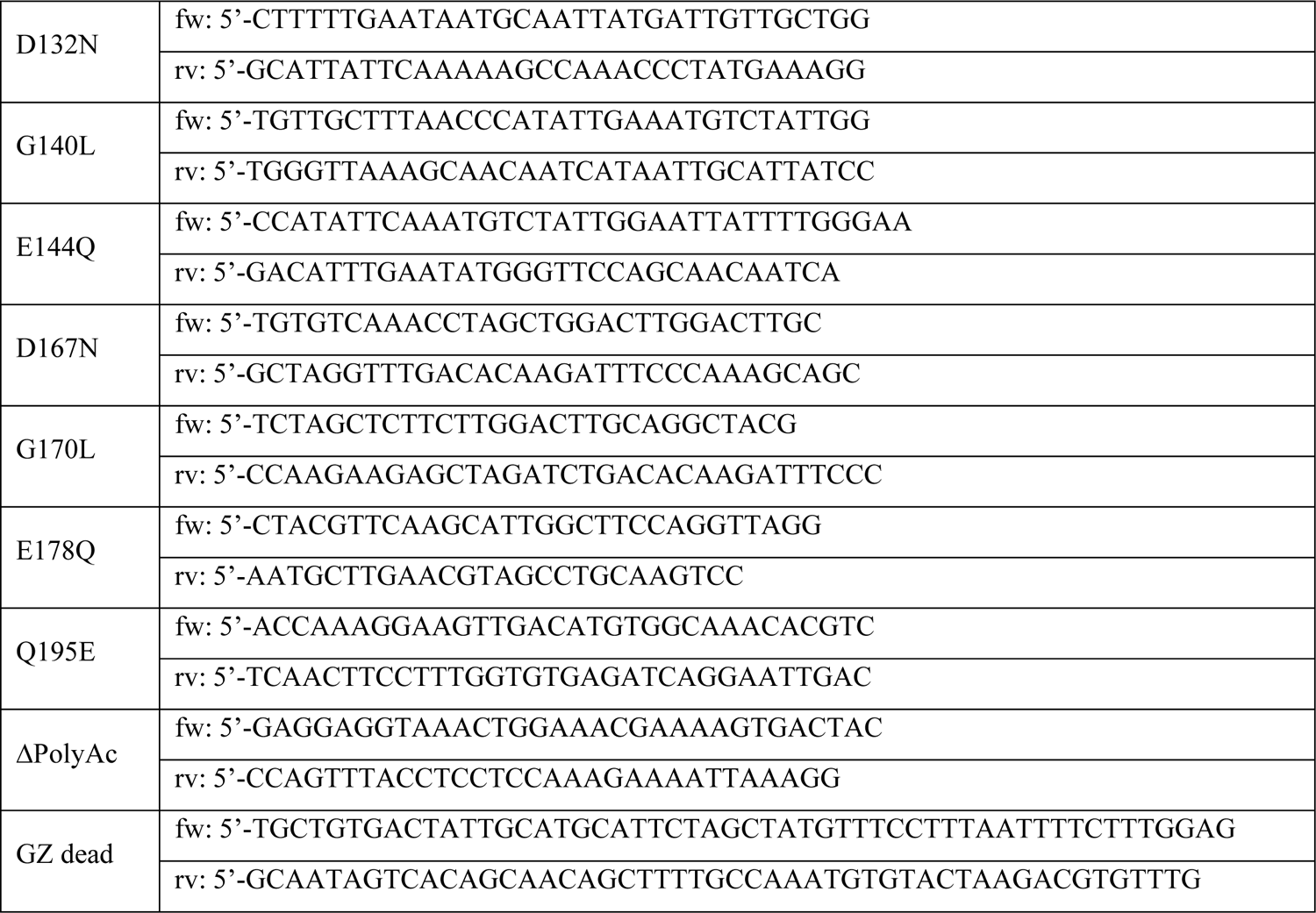

All constructs were verified by Sanger sequencing.

The following siRNAs were purchased from Sigma-Aldrich: esiRNA-RLuc (cat no EHURLUC), esiRNA-TMEM65 (cat no EHU042431), siRNA Universal Negative Control #2 (cat no SIC002), siRNA-TMEM65#1 (cat no SASI_Hs01_00147974), siRNA-TMEM65#2 (cat no SASI_Hs02_00371420), siRNA-TMEM65#3 (cat no SASI_Hs01_00147977), siRNA-TMEM65#4 (SASI_Hs01_00147978).

### Antibodies, SDS-PAGE and Western blot

Cells were lysed in RIPA buffer (150 mM NaCl, 25 mM Tris-Cl pH 8, 1 mM EGTA-Tris, 1% Triton X-100, 0.5% sodium deoxycholate and 0.1% SDS) supplemented with Complete EDTA-free protease inhibitor mixture (Roche Applied Science) and PhosStop (Roche Applied Science) for 30 minutes on ice. Crude extracts were centrifuged 15000 *g* for 10 minutes to remove debris, and proteins in the supernatant were quantified using the BCA Protein Assay Kit (Pierce). Approximately 30 μg of proteins were dissolved in LDS sample buffer (Life Technologies) supplemented with 10 mM dithiothreitol, heated for 5 minutes at 95°C and loaded on 4-12% Bis-Tris NuPage gels (Thermo Fisher Scientific). After electrophoretic separation, proteins were transferred onto nitrocellulose membranes by semidry transfer (BioRad). Membranes were blocked for 1 hour at RT with 5% non-fat dry milk in TBS-T (50 mM Trizma, 150 mM NaCl and 0.1% Tween) and probed with the indicated primary antibodies over night at 4°C. Isotype matched, horseradish peroxidase-conjugated secondary antibodies (1:10000, BioRad) were used followed by detection by chemiluminescence (SuperSignal Pico, Pierce). The following primary antibodies were used: anti-GRP75 (1:1000, SCBT sc-133137), anti-TMEM65 (1:1000, Thermo Fisher Scientific PA5-112762), anti-Flag (1:1000, Cell Signaling #2368), anti SDHA (1:1000, Cell Signaling #5839), anti-PDHA1 (1:1000, Cell Signaling #2784), and anti-MITOK (1:10000, Sigma HPA010980). Western blots are representative of at least three independent preparations. For Western blot analyses of *C. elegans* samples, adult animals were collected in water and snap-frozen in liquid nitrogen. *C. elegans* pellets were then sonicated in RIPA buffer supplemented with protease and phosphatase inhibitors (Roche). Lysates were centrifuged as described above and protein concentrations were quantified via Bradford assay (Sigma-Aldrich). Approximately 20-30 μg of protein was resolved on 8% polyacrylamide gels and transferred onto nitrocellulose membranes using semi-dry Trans-Blot Turbo^TM^ (Bio-Rad). Following antibody incubation (anti-GFP, 1:1000, Roche #118144600001), membranes were developed in ECL and imaged using the Chemidoc imaging system (Bio-Rad). Uncropped images of Western blots used for the assembly of final figures are provided as Supplementary Figures.

### Bioinformatic analyses

To evaluate conservation profiles of MCU and TMEM65, reference proteomes of eukaryotic species were retrieved from Uniprot as fasta files (Release 2022_02, 25-May-2022). Only species containing at least 3000 entries in main (canonical) set were considered for further analysis. Homology searches were performed using phmmer (HMMER version 3.3.2)^1^. Human MCU and TMEM65 protein sequences were used as input against reference proteomes using BLOSUM62 as scoring matrix. Hits with an E-value below 0.05 were considered as homologs. Multiple sequence alignments (ClustalW) and phylogenetic trees (RAxML) were performed using MegAlign Pro (DNASTAR version 17.3.1). Visualization of protein topology was generated using Protter^2^. Prediction of transmembrane domains was performed using TMHMM^3^. The presence of mitochondrial targeting peptides was evaluated using TPP3^4^, MitoFates^5^ and TargetP^6^. To evaluate tissue expression of TMEM65, RNA-seq expression values (TPM) were retrieved from GTEx portal^7^. To normalize to mitochondrial abundance, we calculated the average expression of a large subset of broadly expressed mitochondrial proteins, and normalized TMEM65 expression to these values.

### Mitochondrial isolation and proteinase K protection assay

Mitochondria were isolated from HeLa cells through differential centrifugation as previously described^8^. Mitoplasts were obtained through osmotic swelling by incubating mitochondrial fraction in 20 mM Tris-Cl pH 7.4 for 20 minutes on ice. An equal amount of mitoplasts was treated with proteinase K (100 µg/ml) at room temperature for the indicated time and the proteolytic reaction was quenched by PMSF addition. Samples were then loaded on SDS-PAGE and processed for Western blot as described above. Results are representative of at least three different proteolytic reactions.

### Immunofluorescence and confocal imaging

HeLa cells expressing TMEM65-3xFlag were grown on 24 mm coverslips until 50% confluence. Cells were then washed with PBS, fixed in 4% formaldehyde for 10 minutes and quenched with 50 mM NH_4_Cl in PBS. Cells were permeabilized for 10 minutes with 0.1% Triton X-100 in PBS and blocked in PBS containing 2% BSA for 1 hour. Cells were then incubated with primary antibodies for 3 hours at room temperature and washed 3 times with 0.1% Triton X-100 in PBS. The following primary antibodies were used: anti-HSP60 (1:100, SCBT sc-59567), anti-Flag (1:100, Cell Signaling #2368), The appropriate isotype matched AlexaFluor-conjugated secondary antibodies (1:500, Thermo Fisher Scientific) were incubated for 1 hour at RT and coverslips were mounted with ProLong Diamond Antifade reagent (Thermo Fisher Scientific). Images were acquired on a Leica TCS-SP5-II-RS-WLL equipped with a 100x, 1.4N.A. Plan-apochromat objective. AlexaFluor488 was excited by the 488 nm laser line and images were collected in the 495-535 nm range. AlexaFluor555 was sequentially excited with the 543 nm laser line and signal was collected in the 555-600 nm range. Pixel size was set below 100 nm to meet the Nyquist criterion. For each image, a z-stack of the whole cell was acquired, with a step size of 130 nm. Images are presented as maximum projections of the whole stack and were generated using the Fiji image processing package based on ImageJ^9^. Images are representative of at least three independent transfections. *In-vivo* confocal imaging of lipid droplets was performed using adult nematodes expressing the *idrIs1[dhs-3p::dhs-3::gfp+ unc-76(+)]* transgene. Animals were paralyzed with 20 μM levamisole, mounted onto agarose pads and imaged using the Zeiss Airyscan 2 LSM900 equipped with a 63x oil immersion objective (Zeiss).

### [Ca^2+^] measurements

For experiments in intact cells using bioluminescent probes, HeLa cells were grown on 96-well plates and cotransfected with an Aequorin based probe^10,11^ together with the indicated constructs. siRNAs were delivered through RNAiMAX-mediated reverse transfection 72 hours before experiments. Plasmids were delivered through PEI-mediated forward transfection 36/48 hours before experiments. Untargeted siRNA (esiRNA-RLuc or universal negative siRNA) and/or pcDNA3.1 were used as controls. For aequorin reconstitution, cells were incubated for 1 hour at 37°C with 5 μM coelenterazine in KRB (Krebs-Ringer modified buffer: 125 mM NaCl, 5 mM KCl, 1 mM Na_3_PO_4_, 1 mM MgSO_4_, 5.5 mM glucose, 20 mM HEPES, pH 7.4) supplemented with 1 mM CaCl_2_. After reconstitution, cells were washed and placed in 70 μl of KRB and luminescence was recorded on a PerkinElmer Envision plate reader equipped with a two-injector unit. Each well was monitored for 50 seconds during which histamine was first injected (final concentration is 100 μM) to activate calcium transients, and then a hypotonic, Ca^2+^-rich, digitonin-containing solution was added to discharge the remaining aequorin pool. Output data were analyzed and calibrated with a custom made macro-enabled Excel workbook. Where indicated, cells were pre-treated with either vehicle (DMSO) or 20 μM CGP-37157 (Tocris Bioscience).

For experiments in permeabilized cells using bioluminescent probes, HeLa cells were grown on 13 mm round glass coverslips at 50-80% confluence and cotransfected with a low-affinity mitochondrial-targeted aequorin-based probe together with the indicated constructs. For aequorin reconstitution, cells were incubated for 1-2 hours at 37 °C with 5 μM coelenterazine in KRB supplemented with 1 mM CaCl_2_ and then transferred to the perfusion chamber of a custom-built luminometer^11,12^. Experiments in permeabilized cells were performed in an intracellular-like buffer (IB: 130 mM KCl, 2 mM K_2_HPO_4_, 5mM glutamic acid, 2 mM malic acid, 1 mM MgCl_2_, 20 mM HEPES, pH 7.0), supplemented with additional 10 mM KCl, 10 mM NaCl or 10 mM LiCl as specified. Plasma membrane was permeabilized by perfusing cells for 1 min with 100 μM digitonin dissolved in IB containing 500 μM EGTA. Mitochondrial Ca^2+^ uptake in permeabilized cells was initiated by perfusing IB containing 5 μM Ca^2+^, and influx was blocked by adding 10 μM ruthenium red (RuR). The experiments were terminated by lysing the cells with 100 µM digitonin in a hypotonic Ca^2+^-rich solution (10 mM CaCl_2_ in H_2_O), thus discharging the remaining aequorin pool. The light signal was collected and calibrated into [Ca^2+^] values by an algorithm based on the Ca^2+^ response curve of aequorin at physiological conditions of pH, [Mg^2+^] and ionic strength^13,14^.

For experiments in intact cells using fluorescent probes, HeLa cells were grown on 24 mm round glass coverslips at 30-50% confluence and cotransfected with mitoGCaMP6 together with the indicated constructs. Cells were transferred to an imaging chamber and incubated in KRB supplemented with 1 mM CaCl_2_. Images were acquired on an Olympus IX-73 microscope equipped with a 40x/1.3 N.A. oil-immersion semiapochromat objective (UPLFLN40XO). Excitation light was selected with a Deltaram V high speed monochromator (Photon Technology International) equipped with a 75W Xenon Arc lamp. MitoGCaMP6 emission was collected through a 525/50 filter (Chroma) mounted on an OptoSpin25 wheel (Cairn research). Images were captured with a Kinetix22 sCMOS camera (Photometrics). The system is controlled by Metamorph 7.10 (Molecular devices) and was assembled by Crisel Instruments. MitoGCaMP6 was used in excitation ratiometric mode, where cells are alternatively illuminated every second at 485 and 410 nm^15^. Typical exposure time was 100 milliseconds. After background correction, 485/410 fluorescence ratio was calculated for each cell. Where indicated, 100 μM histamine was added, followed by addition of 3 μM ionomycin and 5 mM CaCl_2_ to check for homogenous probe saturation. Analysis was performed with the Fiji distribution of ImageJ^9^. Data are presented as fluorescence ratio (R, 485/410 nm), and each data point represents one coverslip (containing approximately 1 to 5 individual cells).

### ΔΨ_m_ measurements

The measurement of mitochondrial membrane potential is based on the distribution of the mitochondrion-selective lipophilic cation dye tetramethylrhodamine, methyl ester (TMRM, Thermo Fisher Scientific). Cells were loaded with 20 nM TMRM for 30 minutes at 37°C and then transferred to the imaging system. Images were acquired on a Zeiss Axiovert 200 microscope equipped with a 40x/1.3 N.A. PlanFluor objective. Excitation was performed with a Deltaram V high speed monochromator (Photon Technology International) equipped with a 75W Xenon Arc lamp. Images were captured with a Evolve 512 Delta EMCCD (Photometrics). The system is controlled by Metamorph 7.5 and was assembled by Crisel Instruments. TMRM excitation was performed at 560 nm and emission was collected through a 590–650 nm bandpass filter. On each coverslips images of 5 random fields were acquired with a fixed 200 milliseconds exposure time. At the end of each experiment, 10μM CCCP was added to collapse ΔΨ_m_. data are presented as average background-corrected fluorescence in resting conditions. Data were obtained from at least three independent preparations. All analyses were performed with the Fiji distribution of ImageJ.

### Caenorhabditis elegans strains and maintenance

Nematodes were grown at 20 °C unless differently stated. *C. elegans* was maintained on nematode growth media (NGM) plates and grown on a bacterial lawn of OP50 *E. coli* following standard culture methods. The following strains were used in this study: wild type N2 (Bristol), BAN524 *idrls1[dhs-3p::dhs-3::gfp+unc-76(+)]*, BAN544 *tag-321(bon103)IV*, BAN545 *C03B8.3(bon104)III*, BAN547 *C03B8.3(bon104)III;tag-321(bon103)IV*, BAN580 *C03B8.3(bon104)III;mcu-1(ju1154)IV;tag-321(bon105)IV*, CZ19982 *mcu-1(ju1154)IV*. BAN615 *C03B8.3(bon104)III;tag-321(bon103)IV;idrls1[dhs-3p::dhs-3::gfp+unc-76(+)].* Some strains were provided by the CGC, which is funded by NIH Office of Research Infrastructure Programs (P40 OD010440).

### CRISPR/Cas9 gene editing

Young adult *C. elegans* hermaphrodites were injected with customized injection mix of Cas9 protein (pre-complexed with sgRNAs), target-specific sgRNAs, target donor homology repair template (ssODN) (InVivo Biosystem, Eugene, Oregon, USA) and a plasmid encoding *myo-2p::GFP*. F1 offspring were screened for GFP expression in the pharynx, while F2 were genotyped using specific oligonucleotides upstream and downstream the expected deletions. DNA sequencing was used to confirm the mutation. Selected positive mutants were backcrossed at least five times with wt N2 males.

### DIC microscopy

Eggs from adult nematodes were extracted via bleaching, washed twice H_2_0 and then transferred onto glass slides. Images were acquired with an EPI-SCOPE1 Apotome microscope equipped with a 63x oil immersion objective (Zeiss).

### Hatching assay

Gravid adult animals grown at 20 °C were collected and bleached with a hypochlorite solution. The resulting eggs were transferred on bacteria-seeded NGM plates and incubated at 20 °C or 25°C. Individual young adults were then transferred on freshly bacteria-seeded plates every other day until no eggs were produced. Hatched progeny was quantified over time (up to 120 hours) and the total number of animals summed together. For each experiment, at least 8 nematodes per genotype were used.

### Lifespan assay

Gravid adult animals grown at 20 °C were collected and bleached using a NaClO/NaOH solution to extract eggs. Synchronized populations were then placed in an incubator at 20 °C or at 25 °C, according to the experimental paradigm. Animals were transferred every 2 days and scored at least every other day for touch-provoked movement until death. Animals that died abnormally (e.g., internal hatching, vulva protrusions) were scored as censored. Survival curves were generated using GraphPad Prism software (GraphPad Software Inc., San Diego, USA).

### Sample preparation, LC-MS/MS measurements and database searching

Young adult wt(N2) and *C03B8.3(bon104);tag-321(bon103)* double mutants were collected in water, spun down and stored at −80°C until they were processed for LC-MS/MS. Samples were lysed in 200 μl Lysis buffer (50 mM HEPES (pH 7.4), 150 mM NaCl, 1 mM EDTA, 1.5 % SDS, 1 mM DTT; supplemented with: 1× protease and phosphatase inhibitor cocktail (ThermoScientific)). Lysis was aided by repeated cycles of sonication in a water bath (6 cycles of 1 min sonication (35 kHz) intermitted by 2 min incubation on ice). Approximately, 20 μg of *C. elegans* protein lysates were reduced and alkylated prior to processing by a modified filter-aided sample preparation (FASP) protocol as previously described^16^. Samples were digested overnight with Trypsin (1:20; in 50 mM ammonium bicarbonate) directly on the filters, at 30 °C and precipitated using an equal volume of 2M KCl for depletion of residual detergents. Tryptic peptides were then cleaned, desalted on C18 stage tips and re-suspended in 20 μl 1% FA for LC-MS analysis. MS runs were performed with at least 3 biological replicates.Tryptic peptides were analyzed on a Dionex Ultimate 3000 RSLC nanosystem coupled to an Orbitrap Exploris 480 MS. Peptides were injected at starting conditions of 95% eluent A (0.1% FA in water) and 5% eluent B (0.1% FA in 80% ACN), with a flow rate of 300 nL/min. They were loaded onto a trap column cartridge (Acclaim PepMap C18, 100Å, 5 mm x 300 μm i.d., #160454, Thermo Scientific) and separated by reversed-phase chromatography on an Acclaim PepMap C18, 100Å, 75 µm X 25 cm (both columns from Thermo Scientific) using a 75 min linear increasing gradient from 5% to 31% of eluent B followed by a 20 min linear increase to 50% eluent B. The mass spectrometer was operated in data dependent and positive ion mode with MS1 spectra recorded at a resolution of 120K, mass scan range of 375−1550, automatic gain control (AGC) target value of 300% (3×10^6^) ions, maxIT of 25 ms, charge state of 2-7, dynamic exclusion of 60 sec with exclusion after 1 time and a mass tolerance of 10 ppm. Precursor ions for MS/MS were selected using a top speed method with a cycle time of 2 sec. A decision tree was used to acquire MS2 spectra with a minimum precursor signal intensity threshold of 3×10^5^ for scan priority one and an intensity range of 1×10^4^ −3×10^5^ for scan priority two. Data dependent MS2 scan settings were as follows: isolation window of 2 m/z, normalized collision energy (NCE) of 30% (High-energy Collision Dissociation (HCD)), 7.5K and 15K resolution, AGC target value of 100% (1×10^5^), maxIT set to 20 and 50 ms, for scan priority one and two, respectively. Full MS data were acquired in the profile mode with fragment spectra recorded in the centroid mode. Raw data files were processed with Proteome Discoverer™ software (v2.5.0.400, Thermo Scientific) using SEQUEST® HT search engine against the Swiss-Prot® Caenorhabditis elegans database (v2022-06-14). Peptides were identified by specifying trypsin as the protease, with up to 2 missed cleavage sites allowed and restricting peptide length between 7 and 30 amino acids. Precursor mass tolerance was set to 10 ppm, and fragment mass tolerance to 0.02 Da MS2. Static modifications were set as carbamidomethylated cysteine, while dynamic modifications included methionine and N-terminal loss of methionine, for all searches. Peptide and protein FDR were set to 1% by the peptide and protein validator nodes in the Consensus workflow. Default settings of individual nodes were used if not otherwise specified. In the Spectrum Selector node, the Lowest Charge State = 2 and Highest Charge State = 6 were used. The INFERYS rescoring node was set to automatic mode and the resulting peptide hits were filtered for maximum 1% FDR using the Percolator algorithm in the Processing workflow. A second stage search was activated to identify semi-tryptic peptides. Both unique and razor peptides were selected for protein quantification. Proteins identified by site, reverse or potential contaminants were filtered out prior to analysis. Label-free quantitative MS analysis of the *C03B8.3(bon104);tag-321(bon103)* double mutants in comparison to the wt(N2) reference was performed.

### Statistical analysis of data

Bar graphs and error bars indicate mean ± s.d., and dots show individual data points. Variance was calculated by one-way or two-way ANOVA, as indicated in the legend. When two groups were compared, statistical significance was evaluated using a Mann-Whitney or Student’s t tests, as indicated in the legend. In the case of multiple comparisons, the Holm-Sidak or Dunnet post hoc tests were used. Where applicable, raw data and exact p-values are indicated in source data tables. All analyses were performed with the Origin Pro 2023, GraphPad Prism 8 or Excel (Microsoft).

