## supplementary material for "TMEM65-dependent Ca^2+^ extrusion safeguards mitochondrial homeostasis"

This document contain:

- Supplementary table 1
- Supplementary figures 1-6

### Supplementary table 1

| Statistical analysis of lifespan assays at 25°C | | | |
|---|---|---|---|
| **Genotypes** | **Dead (Censored)** | **Median ± S.E.M** | **Replicates** |
| wt (N2) | 223(57) | 15.00± 0.41 | 4 |
| *tag-321(bon103)* | 185(55) | 13.67±0.33 | 3 |
| *C03B8.3(bon104)* | 216(24) | 13.67±0.33 | 3 |
| *C03B8.3(bon104);tag-321(bon103)* | 296(63) | 13.50±0.29 | 4 |

**Extended lifespan table with reported individual assays**

| Lifespan assay (25°C) | | | | | |
|---|---|---|---|---|---|
| **Exp** | **Genotypes** | **Dead (Censored)** | **Median** | **Max** | ***p* value (compared to wt)** |
| 1 | wt (N2) | 88(17) | 14 | 18 | - |
| | *tag-321(bon103)* | 74(31) | 14 | 17 | 0.1946 |
| | *C03B8.3(bon104)* | 91(14) | 14 | 18 | 0.6802 |
| | *C03B8.3(bon104);tag-321(bon103)* | 119(21) | 14 | 18 | 0.0886 |
| 2 | wt (N2) | 55(20) | 15 | 19 | - |
| | *tag-321(bon103)* | 62(13) | 14 | 19 | 0.1356 |
| | *C03B8.3(bon104)* | 69(6) | 14 | 19 | 0.6986 |
| | *C03B8.3(bon104);tag-321(bon103)* | 53(22) | 13 | 18 | 0.0031 |
| 3 | wt (N2) | 34(6) | 16 | 20 | - |
| | *C03B8.3(bon104);tag-321(bon103)* | 57(12) | 14 | 20 | 0.0352 |
| 4 | wt (N2) | 46(14) | 15 | 20 | - |
| | *tag-321(bon103)* | 49(11) | 13 | 18 | 0.0010 |
| | *C03B8.3(bon104)* | 56(4) | 13 | 17 | 0.0002 |
| | *C03B8.3(bon104);tag-321(bon103)* | 67(8) | 13 | 17 | <0.0001 |

| Lifespan assay (20°C) | | | | | |
|---|---|---|---|---|---|
| **Exp** | **Genotypes** | **Dead (Censored)** | **Median** | **Max** | ***p* value (compared to wt)** |
| 1 | wt (N2) | 72(18) | 25 | 31 | - |
| | *tag-321(bon103)* | 127(32) | 23 | 31 | 0.2118 |
| | *C03B8.3(bon104)* | 133(30) | 23 | 31 | <0.0002 |
| | *C03B8.3(bon104);tag-321(bon103)* | 71(24) | 24 | 30 | 0.1693 |

Figure 1c

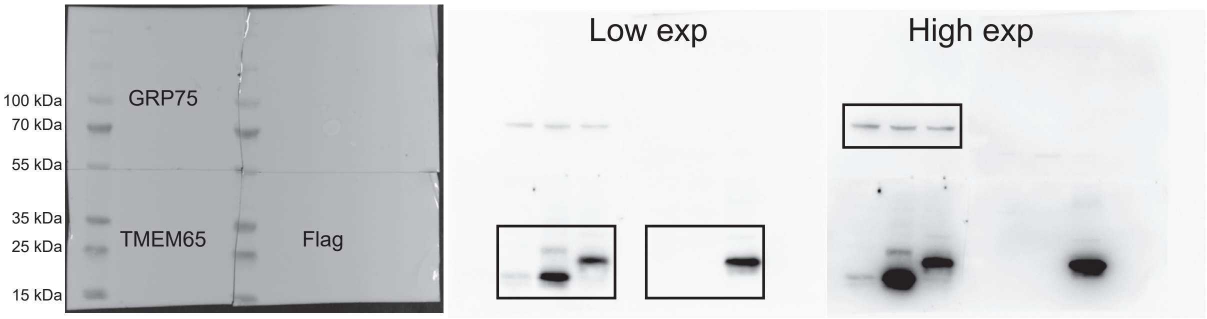


Figure 1e

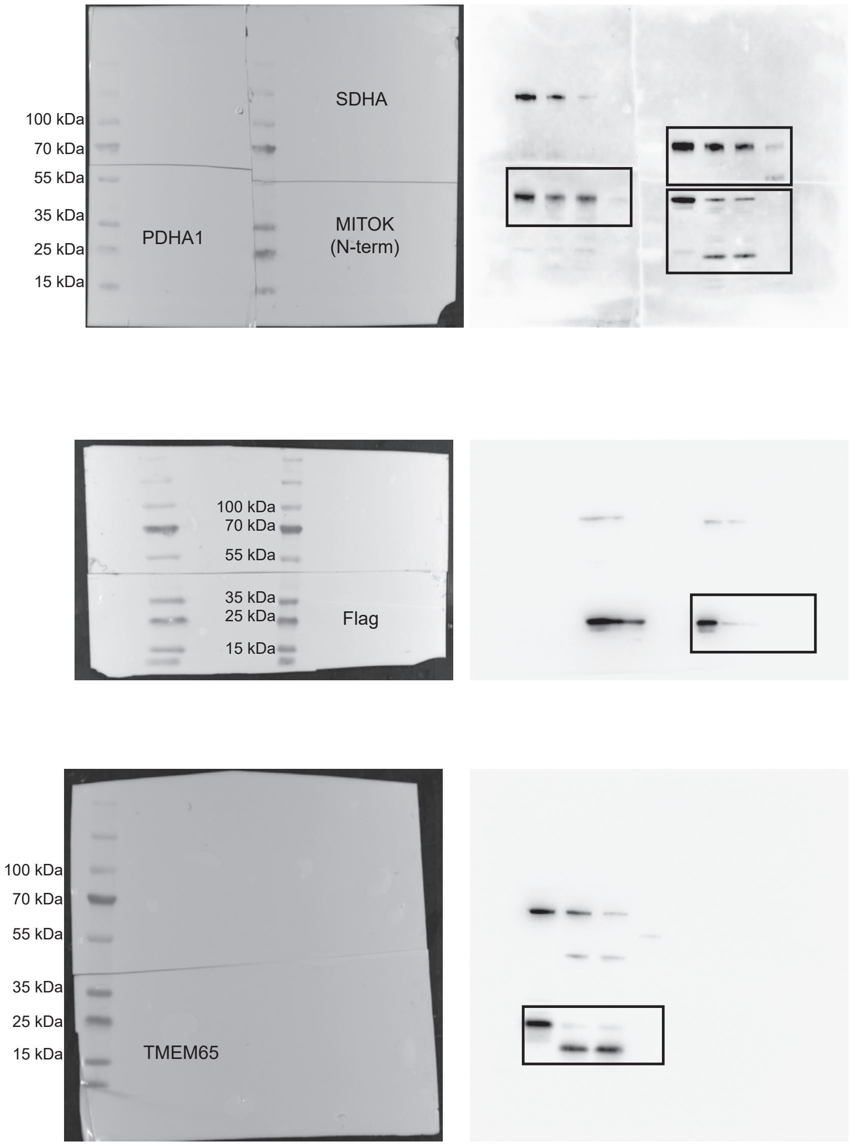


Supplementary figure 1

Extended Data Figure 2e

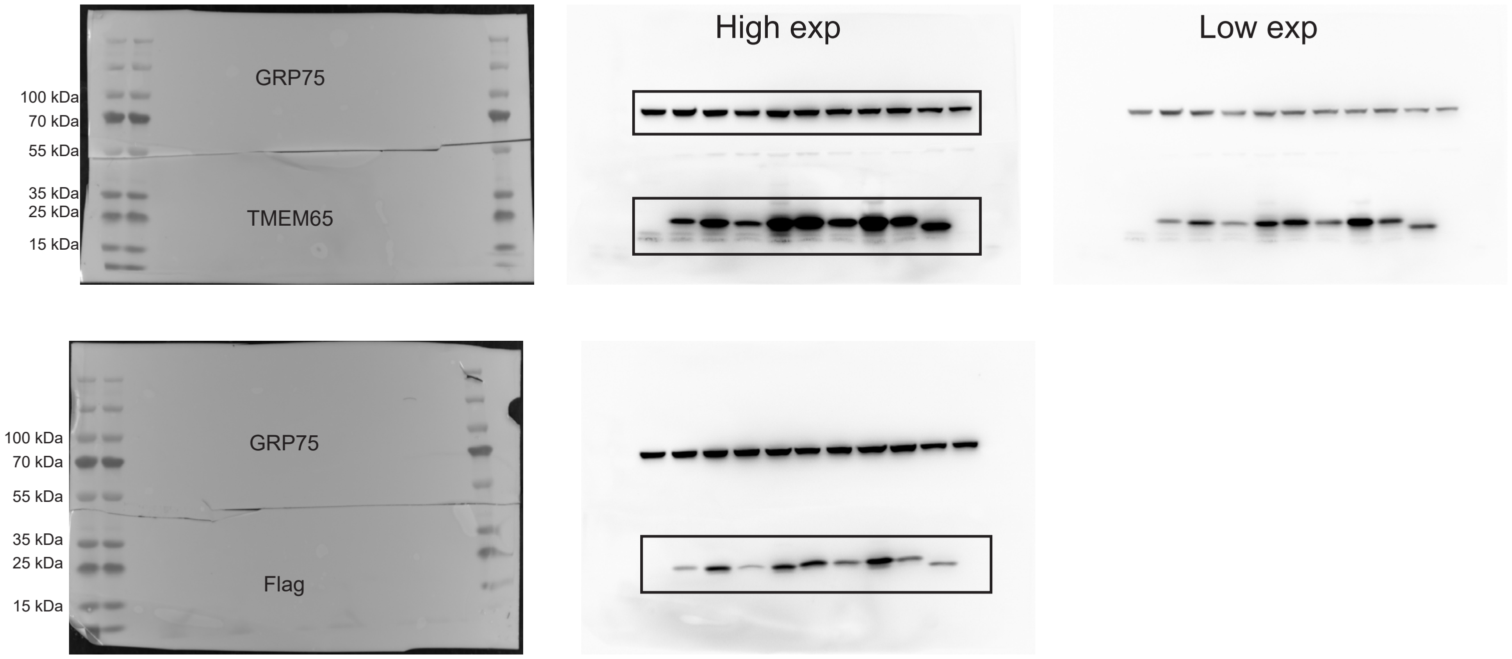


Supplementary figure 2

Extended Data Figure 3e

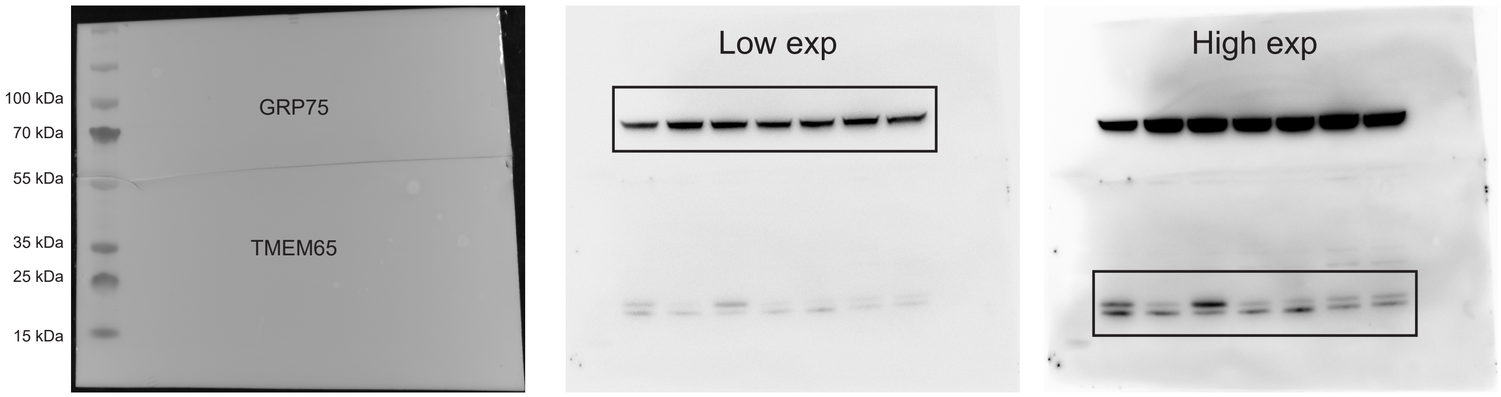


Supplementary figure 3

Extended Data Figure 5d

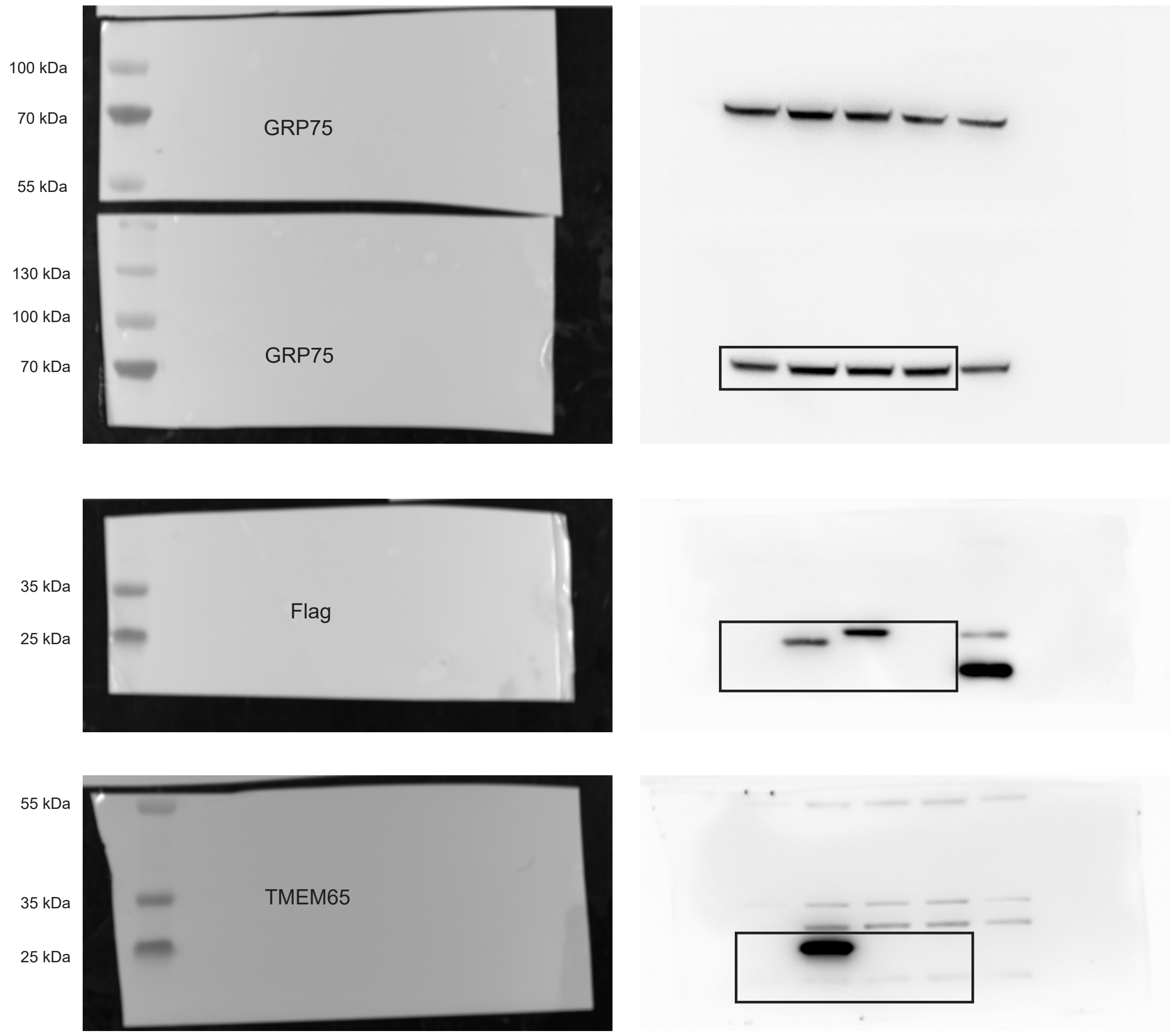


Supplementary figure 4

Figure 4c

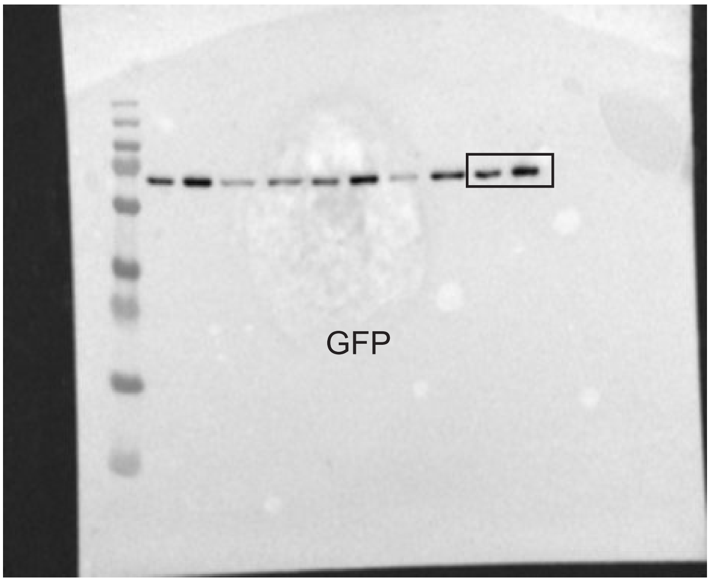


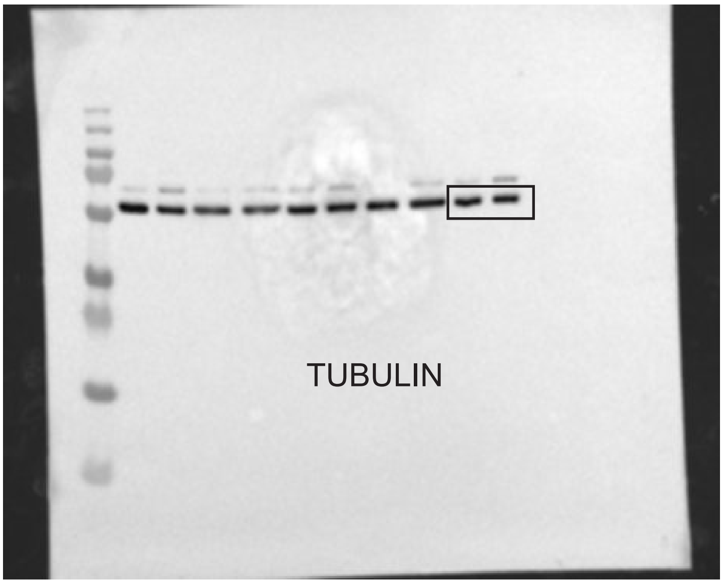


Supplementary figure 5

Extended Data Fig 6d

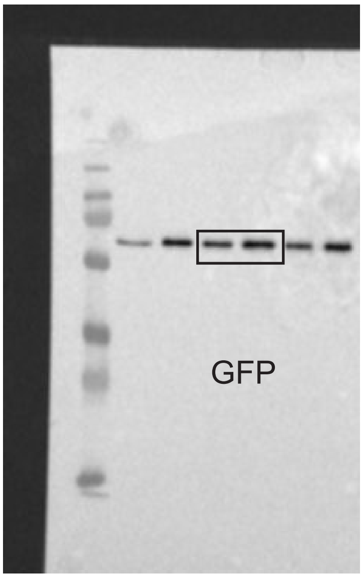


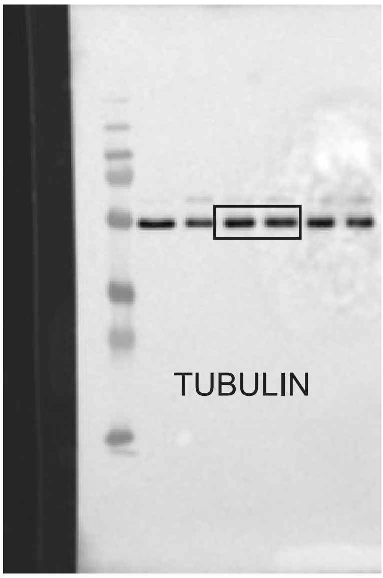
